# Accurate trajectory inference in time-series spatial transcriptomics with structurally-constrained optimal transport

**DOI:** 10.1101/2025.03.19.644194

**Authors:** John P Bryan, Samouil L Farhi, Brian Cleary

**Affiliations:** Faculty of Computing and Data Sciences, Boston University, Boston, MA; Spatial Technology Platform, Broad Institute of MIT and Harvard, Cambridge, MA; Department of Biology, Boston University, Boston, MA; Department of Biomedical Engineering, Boston University, Boston, MA; Program in Bioinformatics, Boston University, Boston, MA; Biological Design Center, Boston University, Boston, MA

## Abstract

New experimental and computational methods use genetic or gene expression observations with single cell resolution to study the relationship between single-cell molecular profiles and developmental trajectories. Most tissues contain spatially contiguous regions that develop as a unit, such as follicles in the ovary, or tubules and glomeruli in the kidney. We find that existing approaches designed to use time series spatial transcriptomics (ST) data produce biologically incoherent trajectories that fail to maintain these structural units over time. We present Spatiotemporal Optimal transport with Contiguous Structures (SOCS), an Optimal Transport-based trajectory inference method for time-series ST that produces trajectory inferences preserving the structural integrity of contiguous biologically meaningful units, along with gene expression similarity and global geometric structure. We show that SOCS produces more plausible trajectory estimates, maintaining the spatial coherence of biological structures across time, enabling more accurate trajectory inference and biological insight than other approaches.

## Introduction

Recent advances in single cell profiling have opened new opportunities to identify developmental trajectories and understand cellular changes underlying dynamic biological processes^1^. Identifying ancestor-descendant relationships allows one to gain insight into the shifts in cell state and cell type composition, and the regulatory networks that guide development, differentiation, and activation. In general, these new technologies have opened two approaches to understand ancestor-descendant relations between cells: prospective genetic fate mapping^2–4^ and computational inference based on high-dimensional single cell profiles^5,6^.

In prospective genetic fate mapping, cells are marked in a way that persists upon cell division (*e*.*g*. with a transgenic fluorescent reporter or a DNA sequence barcode) such that it is possible to identify cells that share a common ancestor, as in technologies such as scGESTALT^7^, LINNEAEUS^8^, and ScarTrace^9^. The advantage of these techniques is that they require minimal inference or assumptions: sets of cells sharing an ancestor can be read out directly. However, with these methods it is not possible to identify and profile the ancestor cell that differentiated into any given set of labeled descendants or to follow the trajectory at intermediate steps. Further, these methods cannot be applied if performing genetic modification prior to sample collection is not an option.

Trajectory methods based on computational inference, on the other hand, do not require manipulation of the sample. For example, RNA velocity methods use observations of “new” versus “old” transcripts to infer the instantaneous vector of change in gene expression^10,11^. Alternatively, pseudotime analysis methods like Diffusion pseudotime (DPT)^12^ or Monocle^13^ are commonly used to understand the dynamics of differentiation trajectories. This family of methods take as input a cell-by-gene count table, and assign each cell (or, in some cases, each cell type) a position on a continuous trajectory. In this way, a more complete estimate can be obtained of cell states all along the trajectory of differentiation. Several methods have recently been developed to incorporate spatial information in obtaining pseudotime labels in individual spatial transcriptomics datasets, including SpaceFlow^14^ and the stLearn package’s pseudo-time-space (PSTS)^15^, enabling researchers to study spatial variability in asynchronous development and cell-cell communication’s effects on differentiation. However, popular pseudotime approaches do not incorporate temporal labels into their analysis of time-series data, which, especially when compounded with the difficulty of integrating multiple datasets due to batch effects, can result in incorrect trajectory inferences.

As an alternative to pseudotime methods in studying time-series transcriptomic data, a family of techniques has been developed that use the mathematical framework of unbalanced optimal transport (OT)^16,17^. OT finds a mapping between two populations that can be interpreted as the ancestor-descendant relationship between cells collected at different time points in the same dynamic process. OT has been applied to time-series scRNA-seq with Waddington-OT (W-OT)^18^, which constrains the mapping so that cells at a first time point (*t*_1_) are mapped to cells with similar gene expression profiles at a second time point (*t*_2_). To apply OT-based trajectory inference to time-series spatial transcriptomics data, methods such as PASTE^19^(and PASTE2^20^, using partial OT), spaTrack^21^, Moscot^22^, and DeST-OT^23^ have been developed, in which, in addition to the gene expression similarity constraint of W-OT, the inference is constrained by pairwise spatial consistency, with the optimization problem assuming that any two cells at *t*_1_ should be about as distant in space as their descendants at *t*_2_. Other spatially-informed methods not based on OT, such as Spateo^24^, have also been developed. While these methods have been shown to capture broad trends of cell type differentiation, the constraints applied by these methods are insufficient to produce biologically accurate results in many biological systems. In particular, nearly every developing tissue is composed in part of spatially contiguous biologically significant units that develop as a unit (*e*.*g*. follicles in the ovary, kidney glomeruli and tubules, small intestine villi, lung alveoli and airways, structures like the cerebellum in the brain, ventricles and atria in the heart, and lobes and lobules in the liver, pancreas, thyroid, and many glands). When OT-based trajectory inference methods only take into account gene expression and global geometry, biologically implausible trajectories are inferred, often splitting these contiguous biological units over time, with cells in a single unit having descendants in multiple units.

Previous work, including the multi-omics data integration method Pamona^25^ have shown that incorporating biological prior assumptions into OT-based methods can improve accuracy. In this paper, we introduce Spatiotemporal Optimal transport with Contiguous Structures (SOCS), which builds on previous methods for OT-based spatiotemporal trajectory estimation by imposing constraints encouraging the coherence of contiguous spatial structures, resulting in estimates where these structures develop as a unit. This enables analysis of the development of structures as a whole, including transcriptomic, morphological, and environmental analysis based on estimated trajectories. We evaluate SOCS by applying it to time-series spatial transcriptomics data from mouse organogenesis (the Mouse Organogenesis Spatiotemporal Transcriptomic Atlas (MOSTA) dataset^26^), and show that trajectory estimates generated by SOCS conform to known differentiation patterns, spatially and in terms of gene expression, while simultaneously preserving meaningful structural units, which other OT-based methods fail to do. We then apply SOCS to time-series MERFISH data collected in mouse ovary after being hyper-stimulated to ovulate, using SOCS to study the maturation and growth of ovarian follicles through ovulation. We confirm that the trajectories obtained by SOCS recapitulate known patterns of follicle maturation, and allow us to make novel biological inferences about the dynamics of biological structures, that are unavailable with existing trajectory inference methods without structural constraints.

## Results

### Spatiotemporal Optimal transport with Contiguous Structures (SOCS)

SOCS creates a mapping between datasets at *t*_1_ and *t*_2_ by solving an optimization problem with constraints based on prior biological knowledge and assumptions about the system. Like other methods, SOCS’s trajectory inference assumes that biological differentiation is relatively efficient, meaning that a cell’s descendants will be relatively similar to its ancestors. As input, SOCS takes spatial transcriptomic datasets acquired at *t*_1_ and *t*_2_ of some dynamic process, and finds a joint distribution over these two datasets, represented by a matrix *T*, in which row *i* of *T* gives the distribution for cell *i* at *t*_1_ of inferred descendants at *t*_2_. *T* is created by solving an optimization problem balancing three requirements: first, that descendants will have more similar gene expression profiles to their ancestors relative to non-ancestors; second, that the geometry of the system will be relatively consistent – that is, the spatial distance between any two pairs of ancestor cells will be similar to the distance between their descendants; and third, that contiguous spatial structures will be preserved over time. This third requirement is the key advantage of SOCS over other spatial OT methods, and is achieved by first defining individual contiguous biological structures (via manual or automatic annotation), then penalizing the objective function when cells belonging to the same structure at *t*_1_ have descendants in different structures at *t*_2_. The approach is illustrated in **Fig. 3a**, and the optimization problem and algorithm are given in detail in **Methods**.

### SOCS Accurately Infers Trajectories in Synthetic Data

Evaluating trajectory-inference methods for spatial transcriptomics data is challenging due to the difficulty of obtaining ground truth trajectories between cells. To begin quantifying the performance of these methods, we used synthetic data. In these synthetic time-series data, *t*_1_ gene expression and spatial coordinates were taken from published ST data^24,26^. For the second timepoint (*t*_2_), we simulated advancing cells’ gene expression in time by adding a scaled version of its RNA velocity (an estimate of the instantaneous change in gene expression), and advancing cells’ spatial coordinates by warping them with spatial transformations.

We considered several scenarios, setting the *t*_2_ spatial coordinates in different ways. First, we warped all spatial coordinates with a single affine spatial transformation. Then, we simulated a structured tissue, in which cells a structure move differently than the surrounding cells. This structured synthetic data is illustrated in **Fig 1b**. We used SOCS, along with other trajectory-inference methods, to infer ancestor-descendant relationships between the *t*_1_ and *t*_2_ datasets, and measured the proportion of cells at *t*_1_ which mapped to the correct descendant at *t*_2_.

**Figure 1.**
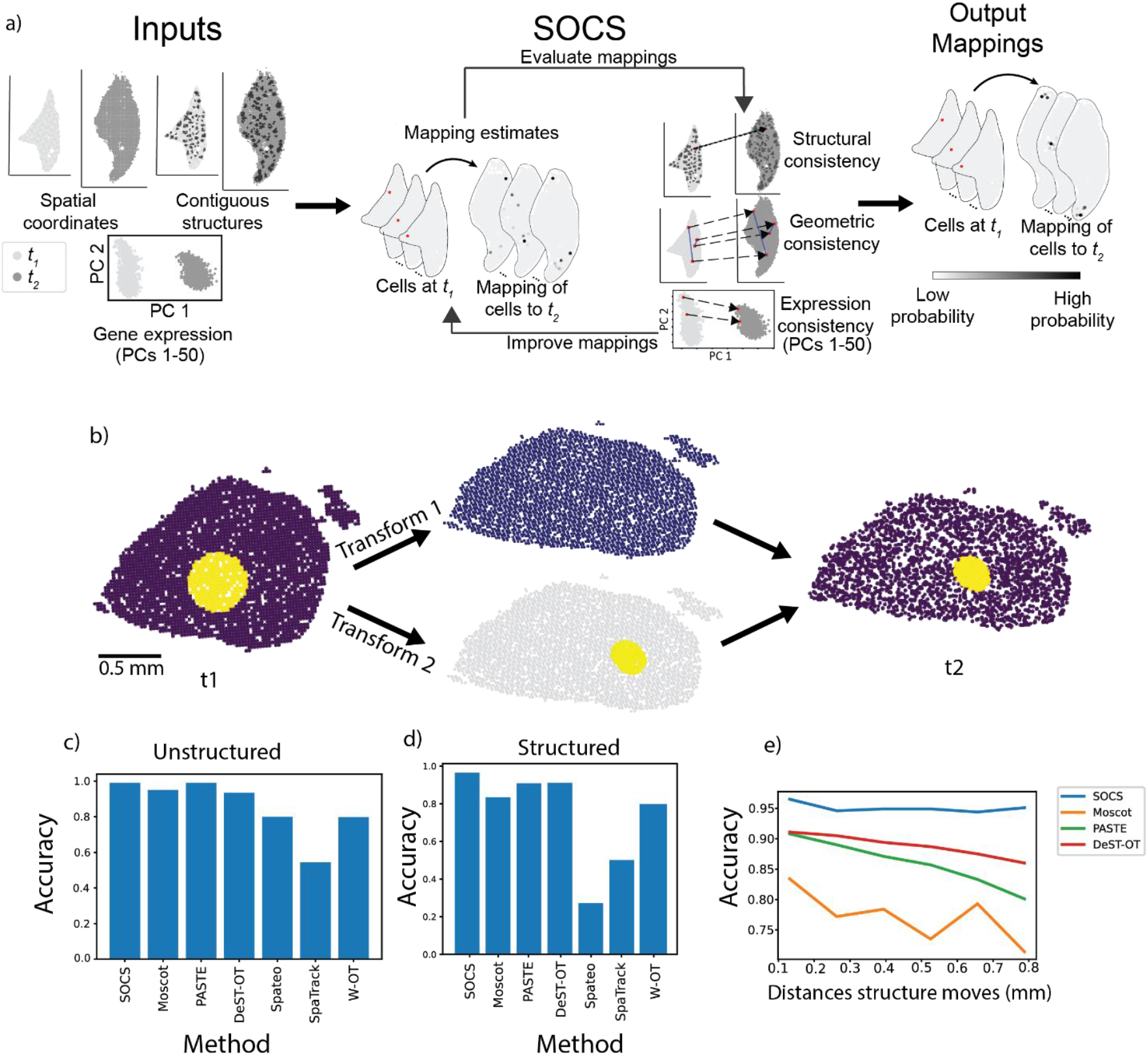
SOCS analysis of synthetic data. **a)** Schematic of SOCS’ operation. Left: inputs to SOCS are spatial coordinates and gene expression data for samples collected at two time points. Center: SOCS minimizes an objective function which encourages consistency in geometry and gene expression. Right: SOCS output is a transport map estimating ancestor-descendant relationships between cells in the two datasets. **b)** Warping of spatial coordinates for synthetic time-series ST data. **c)** Proportion of cells mapping to ground-truth descendant in unstructured synthetic data, by SOCS and other lineage-inference methods. **d)** Proportion of cells mapping to ground-truth descendant in structured synthetic data, by SOCS and other lineage-inference methods. **e)** Variation of proportion of cells mapping to ground-truth descendants as the spatial structure is translated further relative to other cells, by SOCS and other methods.

In the datasets without spatial structure, SOCS performed comparably to the best alternative trajectory-inference methods (**Fig. 1c**). This was expected, as the objective function of SOCS is similar to that of other OT-based approaches when the sample does not include structures. When spatial structures develop independently, SOCS achieved significantly higher accuracy in identifying ancestor-descendant relationships (**Fig. 1d**). As the structure was translated further in relation to the rest of the tissue, the gap between the accuracy of SOCS’s mappings and those produced by other methods grew (**Fig. 1e**). So, SOCS will be particularly valuable in cases where a coherent substructure changes more significantly than its surroundings.

The existence of ground truth in the synthetic datasets allowed us to study the capabilities of SOCS while varying several dimensions. First, when structural boundaries are less clear, structures may be mis-segmented, with cells not actually part of a structure being spuriously attached in the prior. By pre-defining structures in the synthetic data, we were able to examine the effects of mis-segmenting structures. In a given pair of *t*_1_ and *t*_2_ datasets, we tested the effect on ancestor-descendent mapping accuracy of mistakenly identifying cells neighboring the structure with the structure itself. As more spurious cells were added to the structure segmentation, mapping accuracy decreased. However, SOCS achieved higher inference accuracy than other methods with up to 13% of spurious structure cells (**Supplemental Figure 1**). Above that threshold, DeST-OT outperforms SOCS and should thus be preferred in cases with significant uncertainty about structural boundaries, users should prefer DeST-OT.

We also tested the effect of parameter tuning on SOCS’s performance by varying the parameters in the OT objective function. As expected, performance varies depending on the value of these parameters but SOCS continued to outperform other methods even with changes from the optimal settings (**Supp. Fig. 2**). We also used the synthetic data to evaluate the effect of the structural constraint in SOCS. It could be that simply assigning a greater relative weight to global geometry in the SOCS objective function, while not adding the structural constraint. We evaluated maps produced by SOCS with no structural constraint, giving different weights to the global geometry. At all weightings, accuracy was below the accuracy obtained by SOCS with the structural constraint (**Supp. Fig. 3**).

**Figure 2.**
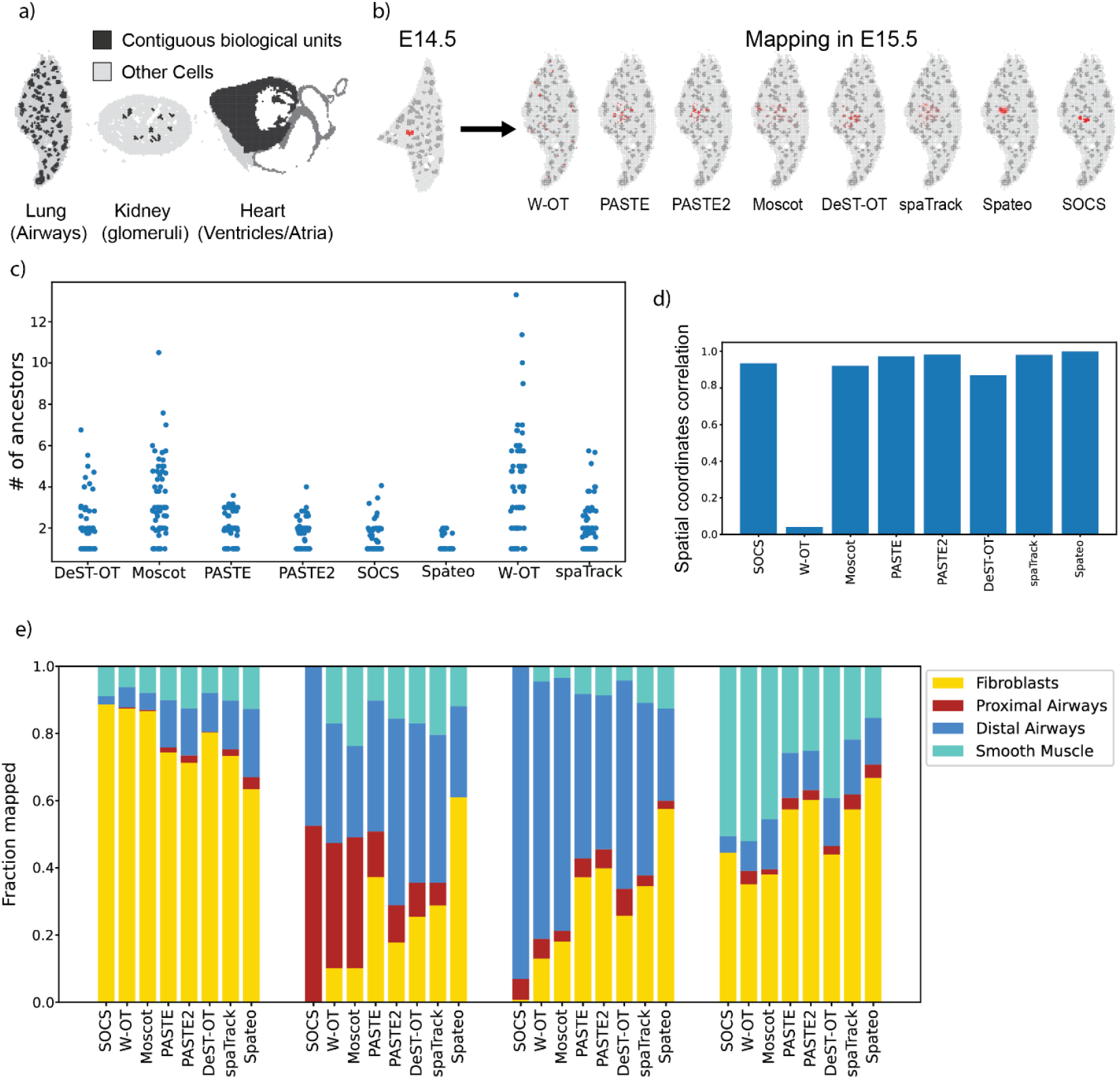
SOCS analysis of time-series Stereo-seq in developing mouse lung: **a)** Examples of contiguous spatial structures in developing mouse organs. **b)** Cells from a single airway cross-section from mouse lung obtained at E14.5 mapped to E15.5 by SOCS and other trajectory-inference methods. **c)** Strip plot showing effective number of ancestor airway cross-sections at E14.5 for each airway cross-section at E15.5, by mapping method. **d)** Pearson correlation between pairwise spatial distances for each cell pair at E14.5 to their mapped pairwise distances at E15.5. Spatially-informed mappings exhibit spatial consistency, while the W-OT mapping does not. **e)** Stacked bar charts giving the proportion of mouse lung cells at E14.5 of each cell type (along the x-axis) mapping to cells of each cell type at E15.5 (represented by the chart’s color), according to the mapping method. SOCS avoids most mapping from epithelial cells (airways) to fibroblasts and smooth muscle cells.

**Figure 3.**
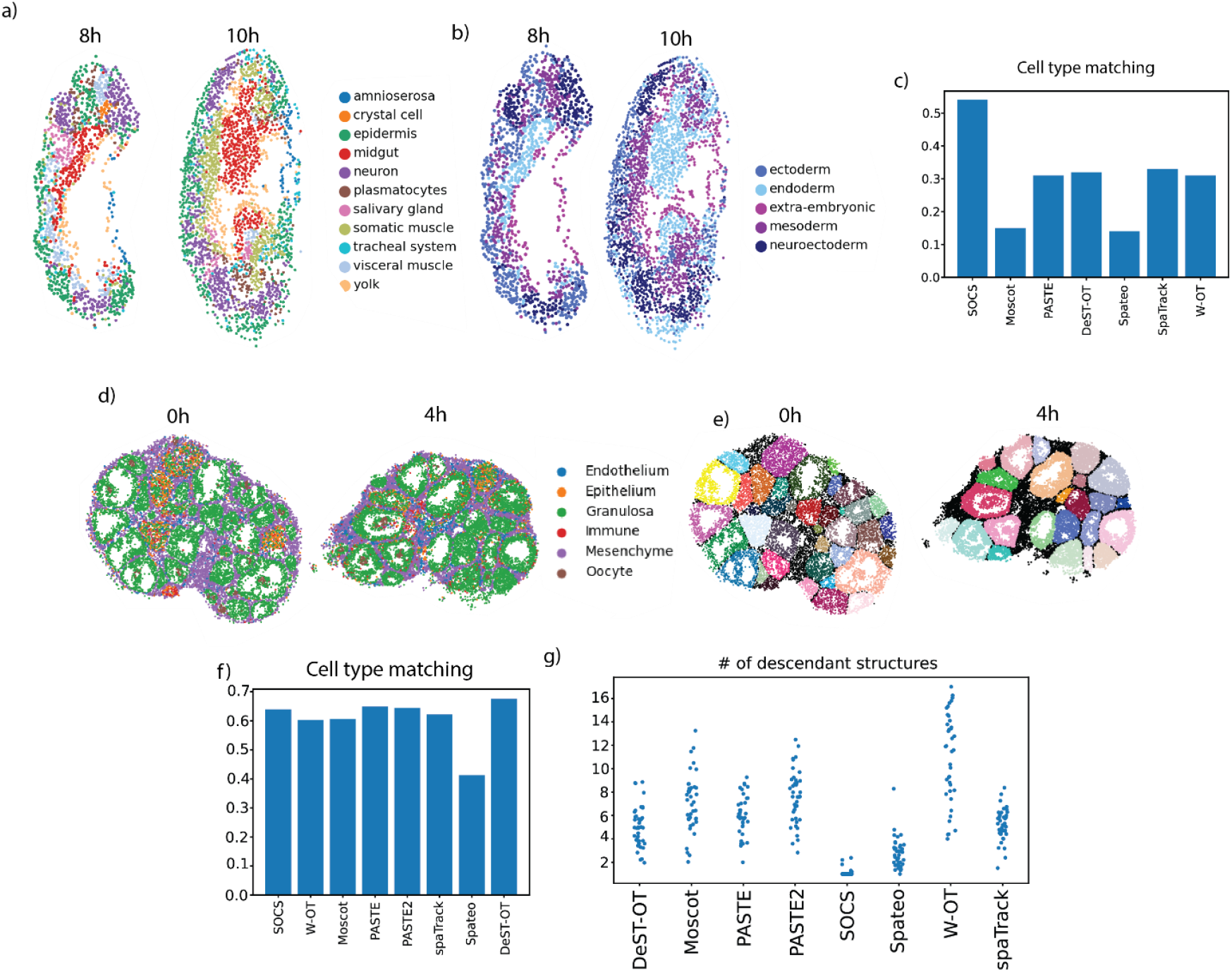
SOCS analysis of drosophila embryogenesis and mouse ovulation. **a)** Slices of Stereo-Seq drosophila embryogenesis datasets from two time-points, colored by annotated cell type. **b)** Slices of Stereo-Seq drosophila embryogenesis datasets from two time-points, colored by annotated germ layer, used as a structural constraint for SOCS. **c)** Proportion of *t*_1_ cells mapping to *t*_2_ cells of the same annotated type, by lineage-inference method. **d)** Slices of Slide-Seq mouse ovulation datasets from two time-points, colored by annotated cell type. **e)** Slices of Slide-Seq mouse ovulation datasets from two time-points, colored by annotated follicle, used as a structural constraint for SOCS. **f)** Proportion of *t*_1_ cells mapping to *t*_2_ cells of the same annotated type in the ovulation data, by lineage-inference method. **g)** Effective number of descendant follicles for each *t*_1_ follicle in the ovulation data, by lineage-inference method.

### SOCS Traces Substructure Development in Developmental ST data

To test SOCS in experimental data, we first used the public mouse organogenesis spatial transcriptomics atlas (MOSTA), which consists of slices of whole mouse embryo, at multiple time points during embryogenesis, profiled with Stereo-Seq^26^, and compared maps between these time points produced by SOCS to maps produced by other methods, W-OT and Moscot. Many developing organs in the MOSTA dataset contain biologically relevant, spatially contiguous structures; we focused on airways in the developing lung, atria and ventricles in the developing heart, and glomeruli in the developing kidney (**Fig. 2a**).

To start, we focused on the developing lung, which has a relatively well-understood developmental trajectory^27^. For our *t*_1_ and *t*_2_ datasets, used the previously annotated lung cells from the whole-embryo MOSTA datasets at E14.5 and E15.5. At these time points the lung is in the pseudoglandular stage, in which the branching architecture of the network of airways is established^28^ and cells are committed to broad cell types. Most cells are epithelial, making up the airways, or mesenchymal, including fibroblasts and smooth muscle cells. We used SOCS to map between the two time points, with the OT problem constrained to maintain the structure of the airway cross-sections and compared the resulting mappings to mappings produced by other computational lineage-inference methods, evaluating the mappings of biological structures, geometry, and cell types.

We first assessed whether the structural constraint of SOCS worked as intended. We qualitatively observed the mapping of individual airways at E14.5 to airways at E15.5, noting that, for the most part SOCS mapped ancestor airways to descendant airways one-to-one, while most other methods mapped any given ancestor airway to many descendant airways (**Fig. 2b**). To quantify this, we computed the effective number of E14.5 airway cross-sections that map to each individual E15.5 airway cross-section, with the expectation that airways might branch, but not merge. We found that in the SOCS mapping, the E15.5 airways (*n* = 77) had a mean of 1.36±0.67 “ancestor” airways, compared to a mean 2.73±2.41 “ancestors” in the W-OT mapping, 2.94±2.30 in the Moscot mapping, 2.71±2.09 in the PASTE mapping, 2.50±1.97 in the PASTE2 mapping, 2.39±1.87 in the DeST-OT mapping, 2.34±1.79 in the spaTrack mapping, and 2.27±1.75 in the Spateo mapping (**Fig. 2c**). Results with 2+ “ancestor” airways run counter to established understanding of lung development. We also examined whether the mappings preserved the geometric relationships between cells (**Fig. 2d**) by comparing the pairwise distances between cells in the E14.5 lung to the pairwise distances between their inferred descendants in the E15.5 lung. In the SOCS mapping, the pairwise distances correlated with the mapped pairwise distances with Pearson’s *r* = 0.93. With the W-OT mapping, the pairwise distances correlated with Pearson’s *r* = 0.03, and with the other spatially-informed trajectory-inference methods the pairwise distances were highly correlated, with Pearson’s *r* between 0.87 and 1.00. Thus, while SOCS and the other spatial methods produce trajectory inferences that preserve global geometry across time, SOCS additionally correctly maps individual contiguous biological structures as a unit, while the other methods fails to map these structures as a unit.

To evaluate the accuracy of the estimated transport maps, we computed the proportion of cells of each type at *t*_1_ (E14.5) mapped to cells of each type at *t*_2_ (E15.5). In the mapping from SOCS, 99.3% of epithelial airway spots in the E14.5 lung mapped to epithelial airways in the E15.5 lung. In contrast, when mapping using all other lineage-inference methods, smaller proportions of epithelial spots mapped to the epithelial airways at E15.5 (**Fig. 2e**). In these under-constrained approaches, when cell types are not well-separated, either in spatial coordinates or gene expression space, cells may map to descendants which are “nearby,” but which are known to be on distinct paths of differentiation. By explicitly defining structures comprised of particular cell types, SOCS avoids this class of error.

We further subdivided epithelial cells into proximal and distal airways, with proximal airways (those airways closer to the “trunk” of the respiratory system) marked by *Sox2*, and distal airways (airways at the branching ends of the respiratory system) marked by expression of *Sox9*^28^. We expect that proximal airways will have descendants that are either proximal airways, or which develop into distal airways as the branching respiratory system grows, and that distal airways, will have only distal airways as descendants. In the SOCS mapping, 49.8% of cells in proximal airways in the E14.5 data mapped to proximal airways in the E15.5 data, with the other 50.2% of cells mapping to distal airways. 92.4% of cells in distal airways in the E14.5 data mapped to distal airways, with 6.8% mapping to proximal airways, with the remaining 0.8% mapping to fibroblasts. In contrast, in the mappings produced by other approaches, there is substantial cross-mapping between epithelial and mesenchymal cells, including both fibroblasts and smooth muscle cells (**Fig. 2e**). Therefore, in methods without structural constraint, minimizing geometric divergence and genetic distance is not sufficient to prevent departure from known lineages.

We confirmed that the improvements seen by SOCS are not limited to the lung by mapping between time-series data in other organs in the MOSTA dataset: heart (**Supp. Fig. 4**) and kidney (**Supp. Fig. 5**). In both datasets, we observe the same patterns as in the lung mappings, with known biological units (glomeruli in kidney and atria/ventricles in heart) splitting in the other methods’ mappings but holding together in the SOCS mapping. This additionally leads to better cell type consistency in the SOCS mapping, without sacrificing spatial consistency. We conclude that the promotion of structural consistency with SOCS produces trajectory inferences that are more biologically accurate than those produced by other OT-based methods.

**Figure 4.**
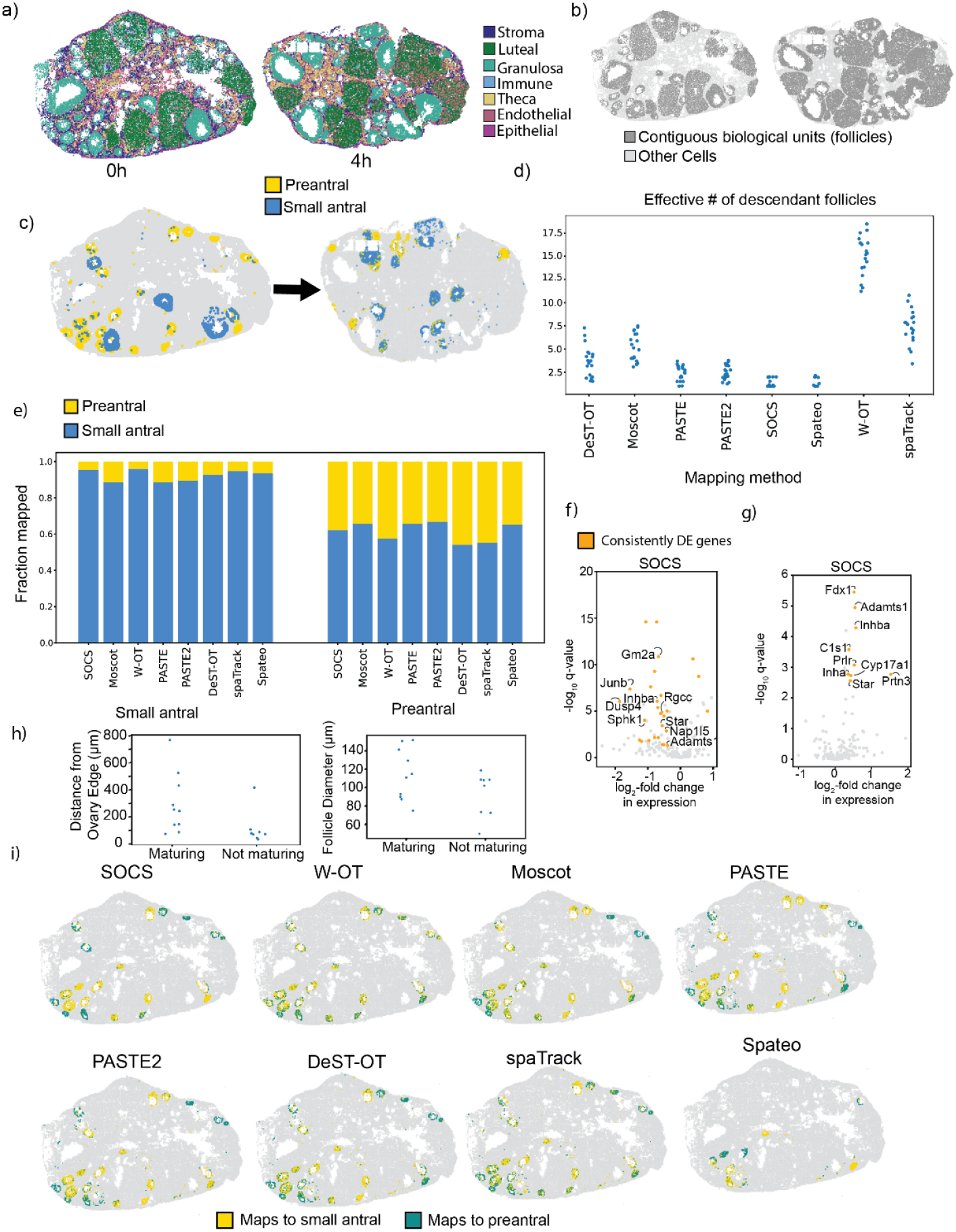
SOCS analysis of time-series MERFISH in mouse ovulation: **a)** Samples of mouse ovary obtained 0 hours and 4 hours after ovulation was stimulated by hCG injection, with cells colored by cell type (legend). **b)** Spatial distribution of follicles, contiguous spatial structures in profiled mouse ovaries. **c)** Immature granulosa cells subdivided by unsupervised leiden clustering into preantral and small antral follicles. **d)** Strip plot showing effective number of descendant follicles at 4h for each mapped follicle at 0h, from trajectory estimation by SOCS and other lineage-inference methods. **e)** Stacked bar charts giving the proportion of small follicle cells of both types at 0h (along the x-axis) mapping to cells of each cell type at 4h (represented by the chart’s color), according to the mapping method. **f)** Volcano plots showing differentially expressed genes (DEGs) in maturing vs. not-maturing preantral follicle cells, from trajectory estimation by SOCS. **g)** Volcano plots showing differential gene expression in stromal cells neighboring maturing vs. not-maturing preantral follicle cells, with consistently DE genes highlighted. **h)** Spatial characteristics of maturing vs. not-maturing follicles. Top: maturing follicles tend to be further from the edge of the ovary. Bottom: maturing follicles tend to have larger diameters. **i)** Spatial distribution of preantral follicle cells at 0h, colored by estimated descendant cell type in 4h, from trajectory estimation by SOCS and other lineage-inference methods.

We wanted to confirm SOCS’s robustness to other types of developmental data. We used SOCS to infer ancestor-descendent relationships in a 3D drosophila embryogenesis dataset (also obtained using Stereo-Seq)^24^ (**Fig. 3a**), and compared to the mappings produced by other datasets. This tested SOCS’s capabilities in 3D ST data, and with data from organisms other than mice. As a structural constraint, we used cells’ annotated germ layer, with the prior assumption that cells belonging to a germ layer at *t*_1_ should have descendants in the same germ layer at *t*_2_. This constraint is slightly different from those implemented in the synthetic data, organogenesis and ovulation datasets: as annotated, these germ layers are not strictly spatially coherent (**Fig. 3b**). As a heuristic to quantify the accuracy of these maps, we computed the proportion of cells at *t*_1_ that mapped to a cell of the same cell type at *t*_2_. We found that SOCS mapped a significantly higher proportion of *t*_1_ cells to cells of the same type than any of the other approaches (**Fig. 3c**). This demonstrates the generalizability of SOCS’s structural constraint approach to other structures which may not be spatially coherent, as well as to three-dimensional data.

Finally, we wished to test SOCS’s robustness to sequencing methodology. We used SOCS to infer ancestor-descendent relationships in a published mouse ovulation dataset obtained using Slide-seq^29^ (**Fig. 3d**). We used ovarian follicles as the structural constraint (**Fig. 3e**), leveraging prior knoweldge that ovarian follicles do not divide during ovulation. We again used the proportion of cells at *t*_1_ mapping to a cell of the same cell type at *t*_2_ as a heuristic for mapping accuracy, and found that SOCS mapped similar proportions of cells at *t*_1_ to cells of the same cell type at *t*_2_ as most other methods (**Fig. 3f**). However, as in the other experimental datasets, we found that SOCS maps cells from a single *t*_1_ follicle to a single *t*_2_ follicle, while the methods not incorporating structural constraints mapped cells from a single *t*_1_ follicle to many *t*_2_ follicles, which is biologically implausible (**Fig. 3g**). This demonstrates that SOCS is robust to changes in sequencing platform.

### SOCS Illuminates Follicle Maturation in Mouse Ovulation

Having evaluated SOCS with synthetic data and sequencing-based time-series ST datasets, we hypothesized that further biological insight could be obtained by studying higher-resolution data obtained using imaging-based ST. We used SOCS to analyze ST data we recently generated in a time course of mouse ovulation using MERFISH^30^—with higher spatial resolution but lower molecular plex than the sequencing-based methods. We used SOCS to map between two mouse ovary datasets at 0h and 4h after hyperstimulation with human chorionic gonadotropin (hCG)^31^ (**Fig. 4a**). In mammalian ovaries, oocytes are surrounded by granulosa cells, forming biological units known as follicles. Through the process of ovulation, certain small follicles (broadly categorized as “preantral” follicles, because they lack the fluid antrum characteristic of more mature follicles) are “recruited,” and mature, with the oocyte and its surrounding follicle growing larger (and acquire a fluid antrum, becoming “antral” follicles), until eventually the follicle ruptures and releases its oocyte for possible fertilization. To study the recruitment and development of these follicles, we identified pre-antral and early antral follicles by first performing unsupervised clustering on gene expression, and then manually annotating clusters by identifying known marker genes^31^. We used SOCS to map between these follicles (**Fig. 4b**), with the OT problem constrained to encourage cells belonging to the same follicle to map as a group (**Fig. 4c**). We also mapped between these datasets with the other before-mentioned lineage-inference methods. As expected, in the SOCS mapping, follicles remained coherent, with follicles at 0h (*n* = 25) having a mean of 1.0±0.45 effective descendant follicles, compared to 14.60±2.38 effective descendant follicles in the W-OT mapping, 2.59±0.79 in the PASTE mapping, 2.69±0.80 in the PASTE2 mapping, 5.53±1.50 in the Moscot mapping, 3.45±1.58 in the DeST-OT mapping, 7.48±2.06 in the spaTrack mapping, and 1.06±0.47 in the Spateo mapping (**Fig. 4d**). SOCS estimated that 38% of cells belonging to preantral follicles at 0h mapped to other preantral follicles at 4h, 62% of cells in preantral follicles at 0h mapped to small antral follicles at 4h; and almost all cells in small antral follicles at 0h mapped to small antral follicles at 4h (95% of small antral follicle cells) (**Fig. 4e**). In the mappings produced by the other methods we saw similar cell-type mapping.

We next compared the population of preantral follicle cells that mapped to preantral follicles (“not maturing”) to the population of preantral follicle cells mapping to small antral follicles (“maturing”). We performed differential gene expression analysis between these populations, and found that 57 genes were differentially expressed in maturing follicles (FDR-adjusted p-value *q* < 0.05, fold change >1.25). For robustness, we also used SOCS to map between the 0h time and a second replicate ovary collected at 4h after hCG stimulation, and repeated the analysis, finding 49 differentially expressed genes, with 28 of the genes being significantly differentially expressed in the same direction in both replicates. Among consistently upregulated genes were *Inhba, Nap1l5, Gm2a, Rgcc, Star*, and *Adamts1*, which are known to be involved with follicle growth and maturation^31^ (**Fig. 4f**), as well as several genes not previously known to be related to this process, including genes such as *Sphk1, Junb*, and *Dusp4*, which were found to be highly differentially expressed (fold change >2.5) with high confidence (*q* < 0.0001), and which may be candidates for future study (**Supplemental Table 3**). When the same analysis was performed with W-OT, the inferred mappings found a similar set of 28 consistently differentially expressed genes, including many of the same genes identified by SOCS, including *Gm2a, Star, Inhba, Adamts1, Dusp4, Junb*, and *Sphk1* (**Supp. Table 4**). It seems reasonable that W-OT might successfully identify differentially expressed genes in cell types categorized by gene expression, given that W-OT takes only gene expression as input. When the same analysis was performed with the other spatially informed inference approaches, greatly varying numbers of differentially expressed genes were found. The inferred mapping by PASTE, Moscot, and Spateo found zero, four, and two consistently differentially expressed genes respectively, while those inferred by PASTE2, DeST-OT, and spaTrack found 18, 35 and 20, respectively. Incorporating global geometry into the optimization problem without including structural information may confound results relating to gene expression, although results may depend strongly on the method used.

In addition to patterns in gene expression, we investigated the morphological characteristics of developing preantral follicles, comparing maturing (*n* = 10) to non-maturing preantral follicles (*n* = 8). Notably, this analysis is facilitated by SOCS but not the other methods, since the latter produce incoherent follicle trajectories that consistently split follicles between descendant types (**Fig. 4g**). We found that maturing follicles had larger mean diameter (114±26 μm vs. 92±23 μm, *p*=0.10 by Welch’s t-test), and were on average further from the edge of the ovary (295±209 μm vs. 112±117 μm, *p*=0.04 by Welch’s t-test) (**Fig. 4h**), which recapitulates known biology – follicles originate at the edge of the ovary and expand and migrate toward the ovary center^32^. We noted that there was significant overlap between the distributions of diameters and distances to the ovary edge, so simply establishing a threshold of diameter and distance to the edge would not be sufficient to identify maturing preantral follicles. We also computed the density of cells in preantral follicles, as it is known that granulosa cells tend to be more densely packed in more mature follicles^33^. We found that maturing follicles were on average slightly denser than not-maturing preantral follicles, but did not find that this difference was statistically significant (0.0082±0.0023 cells/μm^2^ vs. 0.0075±0.0016 cells/μm^2^, p=0.51 by Welch’s t-test) (**Supp. Fig. 7**). Since follicles do not map in a one-to-one manner in W-OT or in the other spatial methods, these kinds of whole-follicle morphological analyses are unique to SOCS.

Finally, to gain insight into the relationship between follicle maturation and the local environment, we studied non-follicular cells neighboring maturing and not-maturing follicular cells. We computed the density of the neighborhoods of maturing and not-maturing preantral follicles, as well as the cell type composition of follicle neighborhoods, but did not find a significant difference in either the neighborhood density (0.0093±0.0039 cells/μm^2^ vs. 0.0107±0.0039 cells/μm^2^, p=0.45 by Welch’s t-test) (**Supp. Fig. 8**) or cell type composition (**Supp. Fig. 9**). However, when we compared the gene expression in non-follicular neighbors of maturing and non-maturing follicles, we found significant differences. We specifically compared the stromal cells within the neighborhoods of these cells, finding 9 genes that were consistently differentially expressed in stromal cells surrounding maturing follicles, including genes known to be expressed in maturing follicles that are now found to be upregulated in neighboring cells, such as *Inhba, Adamts1*, and *Star* (*q* < 0.05, fold change >1.25, **Supp. Table 5**). With the other OT methods, no significantly differentially expressed genes were found (**Fig. 4g**). Of the genes identified by SOCS as differentially expressed, *Prlr* was not previously known to be involved in follicle growth, although it had been identified as relevant to ovarian function and fertility^34^. Given the involvement of *Prlr* in the JAK-STAT signaling pathway, it is possible that it contributes to the induction of cell division in follicle maturation. Thus, in addition to reproducing known follicle biology, SOCS’s structural consistency allows us to form novel hypotheses about the neighborhoods of follicles based on their trajectories.

Overall, by using SOCS to infer differentiation trajectories in mouse ovulation, we were able to find and study populations of maturing and not-maturing preantral follicles. We were able to recapitulate known biological results, identifying differentially expressed genes known to be involved in follicle maturation, as well as morphological differences associated with maturation. We were also able to find previously unreported differentially expressed genes in maturing follicles and their neighborhoods, opening the door to further investigation. Largely due to their lack of constraint on follicle coherence through time, the maps inferred by the other mapping methods were unable to obtain these results.

## Discussion

We have presented SOCS, a method for estimating differentiation trajectories in ST datasets with contiguous spatial biological structures. We have shown that SOCS functions well with different ST technologies, both sequencing-based (with higher-resolution Stereo-Seq and lower-resolution Slide-seq) and imaging-based (MERFISH), showing that SOCS produces trajectory inferences that recapitulate known biological results more effectively than other OT methods, and identifies potentially novel biological results. We suggest that researchers use SOCS to study spatial, genetic, and structural patterns in cellular differentiation, in particular in spatial transcriptomic data sampled over a time-course, where prospective lineage tracing and pseudotime-based approaches will be less suitable.

Accurate cell-level estimates of differentiation trajectories produced by SOCS will enable further analysis and deeper understanding of dynamic biological processes. In addition to organogenesis and homeostatic processes like ovulation, SOCS can be used to study disease progression, in cases like tumor growth and proliferation, or the progress of degenerative diseases. Maps produced by SOCS can be used to nominate molecular drivers of differentiation or disease progression, which could be used to hypothesize therapeutic options. These maps can also be used to identify novel morphological indicators of differentiation or disease, which could eventually be implemented in diagnostic settings.

Incorporating prior biological knowledge into optimization-based methods, as we do with spatial structures in SOCS, should be more generally applicable in OT-based approaches. For example, in Pamona^25^, the authors show that incorporating cell type annotations into an OT-based approach to multi-omics data integration improves the quality of alignment. In general, prior knowledge can be incorporated by increasing the OT cost of associating cells that are known not to be associated, whether because of cell type, spatial structure, lineage, or other prior knowledge.

While we have shown that SOCS is able to accurately estimate differentiation trajectories, and obtain useful biological insights, there are certain limitations to the method. SOCS, as other OT-based techniques, is vulnerable in cases where the time-series samples have strong imbalances in cell type composition. In these cases, the assumption that almost all cells at time point 1 should have at least one descendant at time point 2, and that almost all cells at time point 2 should have an ancestor from time point 1 may cause SOCS to produce results that include artifacts not reflective of actual differentiation patterns. Future efforts may apply principles of partial optimal transport (as in PASTE2^20^), which can further address this limitation. Not all spatiotemporal ST samples are well-suited for SOCS’s structural method. Some biological processes involve spatial structures which do not stay together over time. For example, in corticogenesis, neurons migrate between cortical layers as the brain matures. In cases such as these, we would not recommend applying the structural constraint.

Compared to other trajectory inference methods, SOCS requires more computation time, when run on a CPU (**Supp. Fig. 10**). However, SOCS can be accelerated significantly by processing on a GPU, using a GPU-based implementation of Fused Unbalanced Gromov-Wasserstein OT^35^ implemented in the SOCS software. SOCS is also limited by sampling rate: if time-series data is obtained during a dynamic process at time points separated by a long period of time, tissue structures and gene expression may change dramatically, making it difficult to infer ancestor-descendant relationships.

In this work, we focus on spatial structures, including repeated substructures like airways and ovarian follicles, and unique structures such as embryonic germ layers. However, the approach we take to incorporating this spatial constraint into the optimization problem can easily be extended to other types of structure in future work. For example, in recently-developed brain chimeroids^35^, cells from multiple patient lines are combined in a single tissue. In inferring lineage trajectories between chimeroid datasets, patient line can be used as a structural constraint, with the assumption that cells from a certain line should have descendants from the same line. Other types of structures, including terminal cell types, or germinal layers in embryogenesis, can be used.

Future work may further improve results by combining SOCS with other lineage estimation approaches – for example, clonal barcoding lineage estimation could be integrated with SOCS by treating cells sharing a barcode as belonging to a structure, similar to the contiguous spatial structures. RNA velocity techniques could also be incorporated, with RNA velocity vectors serving as a prior on the mapping of cells in gene expression space. We believe that SOCS will be a useful tool in understanding dynamic spatiotemporal biological processes, including disease progression, and we encourage its adoption in future studies examining time-series ST data.

## Data and code availability

Software implementation of SOCS (including code used to generate synthetic data), together with the data and code used to produce the figures in this paper, are available at https://github.com/algo-bio-lab/SOCS.

## Acknowledgements

We thank Britt Goods, Ruixu Huang, Caroline Kratka, and Jeffery Pea for helpful discussions. We also thank members of the Farhi and Cleary groups for useful feedback. This work was supported by RF1 MH121289 and U01 MH130962 from the National Institutes of Mental Health.

## Author contributions

All three authors conceived of the study. JPB developed and tested SOCS and wrote the first version of the manuscript. BC and SLF supervised the project and edited the manuscript.

## Methods

### Data Acquisition

#### Stereo-seq

For the Mouse organogenesis data analysis, time series data was acquired from the Mouse Organogenesis Spatial Transcriptomics Atlas (MOSTA https://db.cngb.org/stomics/mosta/download/). The obtained data was loaded into anndata^36^ data structures, and subdivided based on the organ annotations provided by the authors. With multiple embryo slices at each time point, we selected slices in which the organs of interest had large cross-sections and in which the biological subunits (airways, atrium/ventricle, glomeruli) were distinguishable through unsupervised gene expression clustering.

For the drosophila embryogenesis data analysis, time-series data was acquired from the Spateo python package^24^. Prior to trajectory inference the slice data was aligned into a 3D dataset by following the steps listed in the Spateo documentation (https://spateo-release.readthedocs.io/en/latest/tutorials/notebooks/4_tdr/3D%20Reconstruction.html).

#### MERFISH

For the MERFISH ovary analysis, data was obtained at multiple timepoints from mice stimulated to ovulated using human chorionic gonadotropin (hCG), as described in^31^, following the standard MERFISH data acquisition protocol using the Vizgen MerScope. Obtained data was saved by the MERSCOPE computational pipeline as cell-by-gene count tables and paired metadata (including spatial coordinates for each cell), and were loaded into anndata data structures. We used those samples with complete ovaries and largely uniform distribution of follicles throughout for our analysis (**Supp. Fig. 5**).

#### Synthetic data

To produce synthetic datasets, we started with scRNA-seq data sampled from mouse dentate gyrus, included as example data in the scVelo^37^ software package. This dataset included precomputed paired RNA velocity vector. We randomly sampled 1000 cells from this dataset and took their gene expression profiles. We assigned each cell a spatial location sourced from a slice from the MOSTA dataset. This paired gene expression and spatial data formed our synthetic data for the first timepoint (*t*_1_). To generate a second timepoint (*t*_2_), we took the *t*_1_ gene expression count table, 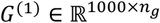 and added the paired RNA velocity vectors, 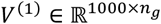, multiplied by a scalar constant *h*:

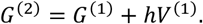

Throughout, we set *h* = 10. To obtain spatial coordinates for the *t*_2_ dataset, we transformed the (*x, y*) coordinates of the *t*_1_ dataset in consistent ways. We produced two types of*t*_2_ datasets: *homogeneous* and *substructured*. For the homogeneous datasets, the *t*_1_ spatial coordinates, 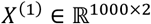are transformed by the application of an affine transform:

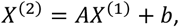

where *A* ∈ ℝ^2×2^ encodes a linear spatial transformation (e.g. scaling, shear mapping, rotation), and *b* ∈ ℝ^2^ encodes a spatial translation.

For substructured datasets, *X*^(2)^ is produced through a multi-step process. First, all spatial coordinates are transformed with a single affine transform:

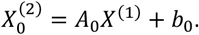

Then, a disc-shaped subset of the *t*_1_ cells’ spatial coordinates, 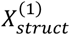is obtained. These spatial coordinates are transformed by a different affine transformation:

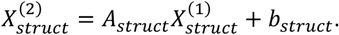

The spatial coordinates in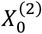are rearranged, with the coordinates in 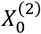originating that overlap with the convex hull of 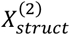 are moved, by mapping the coordinates originating from*t*_1_ cells outside the disc-shaped structure to the set of coordinates in 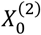that do not overlap with the convex hull of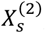 using optimal transport, forming a set of coordinates 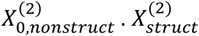 and 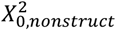 are concatenated, forming *X*^(2)^. This process is illustrated in **Fig. 1a.**

#### Pre-processing

In both the Stereo-seq and MERFISH datasets, we applied standard pre-processing steps with scanpy. After obtaining cell-by-gene count tables, we first filtered out cells with fewer than 10 transcripts. We then normalized counts such that each cell had the same number of counts (normalized to the median total counts in the un-normalized count table), and applied a log-plus-one-transform to the normalized count table.

### Spatial Optimal transport with Contiguous Structures (SOCS)

SOCS takes as inputs ST datasets obtained at two timepoints, *t*_1_ and *t*_2_, which can be represented by cell-by-gene count tables 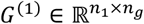 and 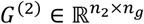, (with *n*_1_, *n*_2_ the number of cells present at*t*_1_ and *t*_2_, respectively, *n*_*g*_ the number of genes profiled, and *G*_*i,g*_ equal to the (normalized and log-transformed) number of mRNA transcripts of gene *g* expressed in cell *i*) paired with vectors giving (x,y) spatial coordinates, 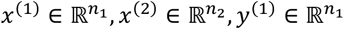, and 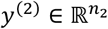. To achieve de-noising and dimensionality reduction, Principal Component Analysis is performed on the count tables *G*^(1)^ and *G*^(2)^, and low-rank representations consisting of the first 50 principal components, 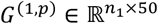 and 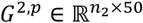are obtained.

SOCS infers the ancestor-descendant relationship of the cells at *t*_1_ and *t*_2_ by creating a map, represented as a matrix 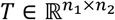 and with element*T*_*i,j*_ representing the “mass” from cell *i* at *t*_1_ transported to cell *j* at *t*_2_ – related to the probability that cell *j* is a descendant of a cell like cell *i*. This is done by solving an optimization problem, minimizing the sum of five arguments:

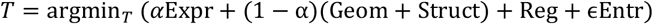

There are five arguments in the objective function, and five tunable parameters: *α, f*_*b*_, *ρ*_1_, *ρ*_2_ and *ϵ*. The first argument is the earth-mover’s distance:

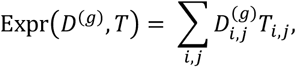

Which imposes a penalty on the gene expression distance:

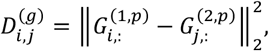

between cells and their descendants/ancestors.

The second is the sum of two Gromov-Wasserstein distances. The first,

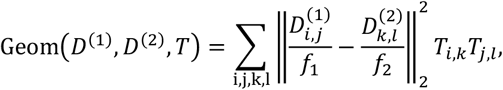

preserves the geometric structure of the sample at *t*_1_ by imposing a penalty on the difference between the distance between two cells at *t*_1_:

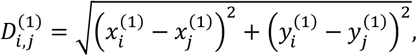

and the distance between their descendants at *t*_2_:

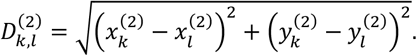

The second:

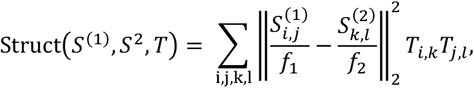

preserves predefined contiguous biological structures by imposing a penalty if two cells belonging to the same structure have descendants belonging to different structures:

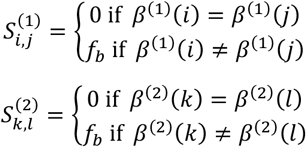

With *β*^(1)^(*x*) and *β*^(2)^(*x*) being maps which assign a cell *x* to a biological structure at *t*_1_ and *t*_2_, respectively. In this paper, these maps were generated from hand-segmentation of the relevant biological structures. We do note that manual segmentation can be time consuming, and introduces an element of human error: automatic segmentation may be useful, especially in cases with many structural elements. We found the SAM3 image segmentation tool to be a flexible, robust approach, and to closely match our manual segmentation in example data (**Supp. Fig. 11**). A tutorial demonstrating the use of SAM3 for this purpose is provided on the SOCS github page.

The parameter *α* trades off geometric and gene expression consistency: if *α* = 0, gene expression is ignored, whereas with *α* = 1, geometry is ignored. To balance the magnitude of (Geom + Struct) and Expr, *D*^(1)^, *D*^(2)^, *S*^(1)^, and *S*^(2)^ are divided by scalar factors *f*_1_ and *f*_2_:

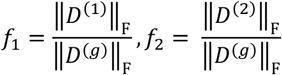

The fourth argument imposes penalties when the number of a cell’s descendants or ancestors departs from the expected number:

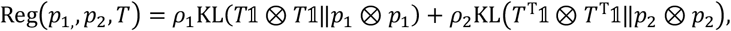

Where *p*_1_ and *p*_2_ are vectors giving the expected number of ancestors and descendants of each given cell, respectively (here set to a uniform distribution), and with the parameters *ρ*_1_ and *ρ*_2_ controlling the magnitude of the penalty for descendants and ancestors, respectively (throughout we found good results from setting *ρ*_1_ = *ρ*_2_ = *ρ*. The final argument:

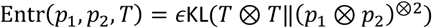

minimizes entropy in the mapping, which has been shown to improve algorithmic convergence^17^. The entropy of the solution *T* can be controlled with the parameter *ϵ*, with the solution being less entropic as *ϵ* increases. We empirically set *ϵ* so that the median Hill number of the rows of *T* was between 1 and 2.

The optimization function is solved with a version of the iterative unbalanced Gromov-Wasserstein algorithm^17^ (the user can provide a maximum number of iterations, *k*, modified to include the arguments Expr(*D*^(*g*)^, *T*) and Struct(*S*^(1)^, *S*^(2)^, *T*).

### Trajectory mapping with other methods

We compared trajectory mappings produced by SOCS to trajectory mappings made by several other lineage-inference methods: Waddington-OT (W-OT), PASTE, Moscot, spaTrack, DeST-OT, Spateo. To produce an estimated transport map with W-OT, we used the `wot` Python package, following the tutorial found at (https://nbviewer.org/github/broadinstitute/wot/blob/master/notebooks/Notebook-2-compute-transport-maps.ipynb). We used the parameters given in the tutorial, with *ϵ* = 0.01, λ_1_ = 1, and λ_2_ = 50. We did not provide an initial estimated growth rate.

To use PASTE and PASTE2, we used the `paste3` Python package, following the tutorial found at (https://raphael-group.github.io/paste3/notebooks/paste_tutorial.html). For PASTE2, we used a mapping fraction of 0.95 for the organogenesis data and 0.99 for the ovulation data.

To produce an estimated transport map with Moscot, we used the `moscot` Python package, following the tutorial found at (https://moscot.readthedocs.io/en/latest/notebooks/tutorials/500_spatiotemporal.html). We ran Moscot with GPU acceleration in a Google Colab notebook.

To make an estimated transport map with Spateo, we used the `spateo` Python package, following the tutorial found at (https://spateo-release.readthedocs.io/en/latest/tutorials/notebooks/3_alignment/1.%20Basic%20usage%20of%20Spateo%20alignment%20for%202D%20slices.html)

To estimate a transport map with DeST-OT, we used the Python package found at (https://github.com/raphael-group/DeST_OT), using the `align` function.

To estimate a transport map with spaTrack, we used the `spaTrack` Python package, using the `get_ot_matrix` function.

### Cell type annotation

We annotated cells according to their type by applying typical single-cell pipeline steps using scanpy. In the Stereo-seq data, we first isolated spots which had been annotated by the MOSTA authors as belonging to the organ of interest. After pre-processing as described above, we performed dimensionality reduction with principal component analysis (PCA), retaining the top 50 principal components (PCs), and constructed a neighborhood graph with the PC representation. To subdivide organs into broad cell types, we performed unsupervised Leiden clustering, using resolution parameters chosen empirically (see **Supp. Table 1** for values chosen). In most cases, we were able to annotate these cell types by visualizing their spatial distribution in the organ. In lung, we annotated cell types by comparing expression levels of known markers of fibroblasts, smooth muscle, and epithelium^27^. We further isolated the spots identified as epithelial cells, and re-clustered to identify proximal and distal airways.

In the MERFISH ovary data, we applied similar pre-processing and clustering steps. After applying the pre-processing steps described above, PCA dimensionality reduction was performed, keeping the top 50 PCs, and a neighborhood graph was generated from the PC representation. To obtain high-level cell types, leiden clustering was performed (see **Supp. Table 2** for resolution parameters chosen). Clusters were annotated by comparison of expression of known marker genes of common ovarian cell types^31^. Granulosa cells were isolated for further subclustering: clusters of cells annotated as granulosa could be divided into two broad groups with leiden clustering on gene expression, with the two groups corresponding to mature, large antral follicles, and smaller, immature follicles. We identified the immature granulosa cells and performed PCA on just those cells, again created a neighborhood graph, and performed Leiden clustering, to obtain subclusters corresponding to preantral follicles and small antral follicles.

### Evaluation Metrics

#### Identifying putative descendants

For certain comparisons (**Fig. 1 d-f, Fig. 2 e-i**), it was desirable to associate each cell at *t*_1_ with a single putative descendant cell at *t*_2_. We did this in a probabilistic fashion: for each cell *i* at *t*_1_, we first obtained its “normalized mapping vector” 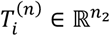, by normalizing the *i*th row of *T* such that its elements sum to 1:

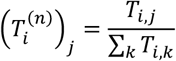

We then used this to define a discrete cumulative distribution function 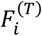 where 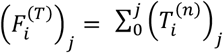. We then sampled from a uniform probability distribution *x*∼*U*_[0,1]_ and identified the descendant cell as *i*′ such that 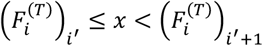.

As the rows of *T* can be interpreted as a discrete probability distribution on the possible*t*_2_ cells, if the maximum value of 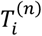is low, this can indicate a low confidence in mapping cell *i*. A threshold on 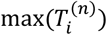 can be applied to filter out *t*_1_ cells with low-confidence mappings prior to downstream analyses.

We found that cells belonging to structures had higher-probability mappings (**Supp. Fig. 12b**), but that thresholding on 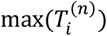 did not result in more accurate maps (**Supp. Fig. 12c**).

#### Evaluating synthetic data results

In the synthetic data, we had established ground-truth ancestor-descendant relationships. To evaluate the accuracy of an obtained lineage inference map, we computed the proportion of cells at *t*_1_ which map to the correct cell at *t*_2_.

#### Evaluating geometric consistency

We evaluated the geometric consistency of the obtained map by comparing each pairwise distance between cells at *t*_1_, 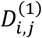to the distance between those cells’ descendants at *t*_2_, 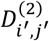. For an aggregated metric, we computed the Pearson correlation between the vectors vec(*D*^(1)^) and vec(*D*^(1)^′), with vec(·) vectorization of a matrix, and *D*^(1)^′ the matrix giving the pairwise distances at *t*_2_ between the putative descendants of the cells at *t*_1_:

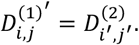

#### Evaluating cell type consistency

As described above, we annotated each dataset with cell type labels, which we can represent as 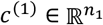 and 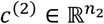, with 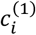giving the cell type label of the*i*th cell at *t*. While not true in all cases, a useful heuristic for accurate mapping is the proportion of cells at *t*_1_ that map to cells of the same type at *t*_2_. For each cell type *k*, we computed 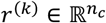, with *n c* the number of unique cell types, and 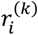giving the percentage of cells of type *k* at *t*_1_ mapping to a cell of type*i* at *t*_2_:

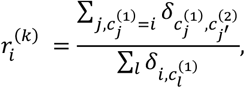

Where *δ*_*i,j*_ is the Kronecker delta.

#### Evaluating structural consistency

As described above, we annotated each dataset with structural labels indicating whether a given cell belongs to a biologically meaningful spatial structure. Similar to the procedure evaluating cell type consistency, for each structure *k*, we computed 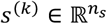, with *ns* the number of unique structures, and 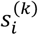giving the percentage of cells belonging to structure*k* at *t*_1_ mapping to a cell of type*i* at *t* :

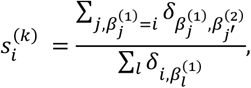

Where *δ*_*i,j*_ is the Kronecker delta. As a measure of the structural consistency of the mapping, we computed the Hill number of each structure-mapping vector *s*^(*k*)^:

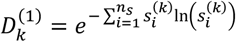

which gives the “effective number” of structures mapped to by structure *k*.

#### Identifying populations with mapping

As described in **Results**, in mappings produced by SOCS, preantral follicle granulosa cells almost entirely mapped either to the same cell type, or the small antral follicle granulosa cell type. We were interested in comparing the population of preantral follicle cells that mapped to the same cell type (not-maturing cells) to the population of preantral follicle cells that mapped to small antral follicles (maturing cells). As described below, we were able to perform differential gene expression on a cell-by-cell level, both for the cells in these populations, and for their neighbors. In the SOCS mapping it was additionally easy to visually identify whole small immature follicles that mature vs. not (**Supp. Fig. 5**). We identified maturing follicles as those with 80% or greater of their cells mapping to maturing follicle cells, and identified not-maturing follicles as those with 80% or greater of their cells mapping to not-maturing follicle cells.

#### Differential expression analysis

To identify genes that are differentially expressed in different populations, such as the maturing vs. not-maturing small immature follicle cells, we used the python package diffxpy^38^ to perform a Wald hypothesis test. For a given test, we identified genes as differentially expressed if the hypothesis test returned an adjusted p-value of 0.05 or lower, and if the fold-change in expression between populations was larger than 1.5. We used this strategy to identify differentially expressed genes in the small immature follicle cells in the 0h dataset, based on their mapping to two replicates of 4h datasets. For robustness, we identified a gene as consistently differentially expressed if the gene was differentially expressed in both tests, and in the same direction (*i*.*e*., upregulated in both or downregulated in both).

#### Morphological characteristics

We were interested in comparing the morphological characteristics of maturing vs. not-maturing small immature follicles. Specifically, we were interested in comparing follicle diameters, and the distance of follicles from the edge of the ovary.

Measurements of follicle diameter can be ambiguous due to the irregular shape of many follicles. To compute the diameter, we first identified the boundaries of the follicle by finding the convex hull of the list of coordinates of the cells making up the follicle. We identified the centroid of each follicle as the centroid of a polygon constructed from the convex hull. We then computed the maximal length a line segment parallel to the x-axis (with the origin at the follicle centroid) spanning the follicle, and repeated this for 180 line segments rotated by evenly spaced (1°, 2°, …, 180°) from the x-axis. We used the mean length of these line segments as our diameter measure. We compared the mean size of the maturing and not-maturing small immature follicles, and tested for significance using a two-sample t-test.

To compute the distance of a follicle from the edge of the ovary, we first manually outlined the border of the ovary using WebPlotDigitizer^39^. For each follicle, we computed the smallest distance between each cell in the follicle and the border of the ovary using the python package shapely, and computed the mean distance for cells in the follicle. We compared the average distance from the edge of the maturing and not-maturing small immature follicles, and tested for significance using a two-sample t-test.

To compute cell density of the follicles, we first computed the area of each follicle (in this case, hand-segmented to account for empty space of the follicle atria), again using the python package shapely. We counted the number of cells contained within the segmented area, and computed the ratio of cells per unit area (**Supp. Fig. 6**). We compared the average density of the maturing and not-maturing preantral follicles, and tested for significance using a two-sample t-test.

#### Neighborhood analysis

We were also interested in the neighbors of granulosa cells. To identify neighborhoods, we first used Delaunay triangulation^40^ to produce a graph representation of the geometry of the ovary, with each node representing a cell. We identified cells as being neighbors if they are separated by a distance less than a set threshold, and their nodes are connected by a single edge in the graph. To compare neighborhoods of different populations, like the maturing vs. not-maturing small immature follicles, we identified all cells neighboring a cell in the population that were not themselves members of the population, isolated neighborhood cells belonging to a cell type of interest, and used the process described above to identify consistently differentially expressed genes.

With the SOCS mapping, in which we were able to easily label whole follicles as maturing and not-maturing, we also characterized the neighborhoods of follicles. Firstly, we found all first- and second-degree neighbors of a follicle’s cells using the method described above. We computed the proportion of the follicle’s neighboring cells that belonging to each high-level cell type. We used a two-sample t-test to compare the average proportion of each cell type in the maturing and not-maturing preantral follicles.

We also computed the density of follicles’ neighborhoods. To do this, we began by again finding all first-and second-degree neighbors of a follicle’s cells. Given these neighboring cells, we found the convex hull of these cells, and computed its area (**Supp. Fig. 7**). From this area we subtracted the area of the segmented follicle to obtain the “neighborhood area.” We divided the number of neighborhood cells by this area to obtain the neighborhood density, and compared the average neighborhood density of maturing and not-maturing preantral follicles, and tested for significance using a two-sample t-test (**Supp. Fig. 8**).

