## Supplementary Material for "Accurate trajectory inference in time-series spatial transcriptomics with structurally-constrained optimal transport"

**Supplemental Materials**

**Supplemental Table 1: Clustering resolution parameters for organogenesis data**

| Dataset | Leiden resolution parameter |
| --- | --- |
| Heart E14.5 | 0.25 |
| Heart E16.5 | 0.35 |
| Kidney E15.5 | 0.45 |
| Kidney E16.5 | 0.85 |
| Lung E14.5 | 0.65 |
| Lung E15.5 | 0.3 |
| Lung Epithelium E14.5 | 0.45 |
| Lung Epithelium E15.5 | 0.45 |

**Supplemental Table 2: Clustering resolution parameters for ovulation data**

| Dataset | Leiden resolution parameter |
| --- | --- |
| 0h Ovary | 1.5 |
| 4h Ovary | 1.5 |
| 0h Immature follicles | 0.15 |
| 4h Immature follicles | 0.15 |

**Supplemental Table 3: Differentially expressed genes in maturing preantral follicles (SOCS map)**

| Gene Name | Fold-change | Log_10_(-q-value) |
| --- | --- | --- |
| Rspo1 | -1.754421 | 4.998933 |
| Wnt6 | -1.457247 | 8.737851 |
| Pcsk6 | -1.290005 | 10.60675 |
| Adamts1 | 1.32581 | 1.306304 |
| Rragd | 1.33831 | 5.012801 |
| Nap1l5 | 1.35748 | 2.877374 |
| Nr4a1 | 1.4304 | 4.481518 |
| Prlr | 1.43665 | 1.404911 |
| Fzd1 | 1.47788 | 1.404911 |
| Star | 1.4885 | 3.440808 |
| Rgcc | 1.49512 | 4.862264 |
| Fdx1 | 1.52156 | 6.683389 |
| Mgarp | 1.53592 | 4.742552 |
| Gm2a | 1.60467 | 10.86491 |
| Runx1 | 1.61162 | 2.155102 |
| Coch | 1.63063 | 5.386973 |
| Inhba | 1.65062 | 6.07295 |
| Apoe | 1.66863 | 14.58734 |
| C1s1 | 1.74246 | 9.269438 |
| Runx2 | 1.74501 | 2.165012 |
| Krt8 | 1.89647 | 7.621121 |
| Vim | 1.96513 | 1.851203 |
| Mro | 2.09918 | 14.58734 |
| Sphk1 | 2.15805 | 4.017297 |
| H2-Ab1 | 2.29504 | 1.75097 |
| Rtp4 | 2.39654 | 1.891041 |
| Junb | 2.93652 | 7.374893 |
| Dusp4 | 3.66005 | 6.030907 |

**Supplemental Table 4: Differentially expressed genes in maturing preantral follicles (W-OT map)**

| Gene name | Fold-change | Log_10_(-q-value) |
| --- | --- | --- |
| Pcsk6 | -1.444813 | 30 |
| Wnt6 | -1.422421 | 7.497008 |
| Rspo1 | -1.408379 | 1.92808 |
| Vcan | -1.339587 | 1.928102 |
| Fkbp6 | -1.323484 | 1.332118 |
| Rasd1 | -1.317828 | 13.69269 |
| Kitl | -1.262368 | 4.842375 |
| Aldh1a1 | 1.31597 | 6.062563 |
| Cyp17a1 | 1.3202 | 1.942498 |
| Cyp11a1 | 1.34254 | 2.293042 |
| Fdx1 | 1.34435 | 3.602534 |
| Gm2a | 1.37122 | 5.126492 |
| Adamts1 | 1.39685 | 1.783297 |
| Apoe | 1.40973 | 6.737819 |
| Inhba | 1.47267 | 3.979453 |
| Nr4a1 | 1.51851 | 6.020553 |
| C1s1 | 1.5338 | 5.905647 |
| Runx2 | 1.55372 | 1.55525 |
| Egr1 | 1.56617 | 2.68166 |
| Mro | 1.56701 | 5.902857 |
| Runx1 | 1.58271 | 2.05766 |
| Coch | 1.59561 | 5.081843 |
| Mgarp | 1.68814 | 6.772438 |
| Star | 1.74612 | 6.354732 |
| Prlr | 1.98017 | 4.042679 |
| Sphk1 | 2.01921 | 3.624881 |
| Dusp4 | 2.07076 | 3.2271 |
| Junb | 2.88375 | 7.578948 |

**Supplemental Table 5: Differentially expressed genes in stromal neighbors of maturing preantral follicles (SOCS map)**

| Gene name | Fold-change | Log_10_(-q-value) |
| --- | --- | --- |
| Prtn3 | 2.892106 | 2.770555 |
| Inhba | 1.48547 | 4.287642 |
| Adamts1 | 1.459067 | 4.947472 |
| Prlr | 1.442901 | 3.073363 |
| Fdx1 | 1.435678 | 5.452425 |
| Cyp17a1 | 1.34844 | 2.730145 |
| Star | 1.337649 | 2.557454 |
| C1s1 | 1.302854 | 3.578336 |
| Inha | 1.280972 | 2.770555 |

**Supplemental Figures**

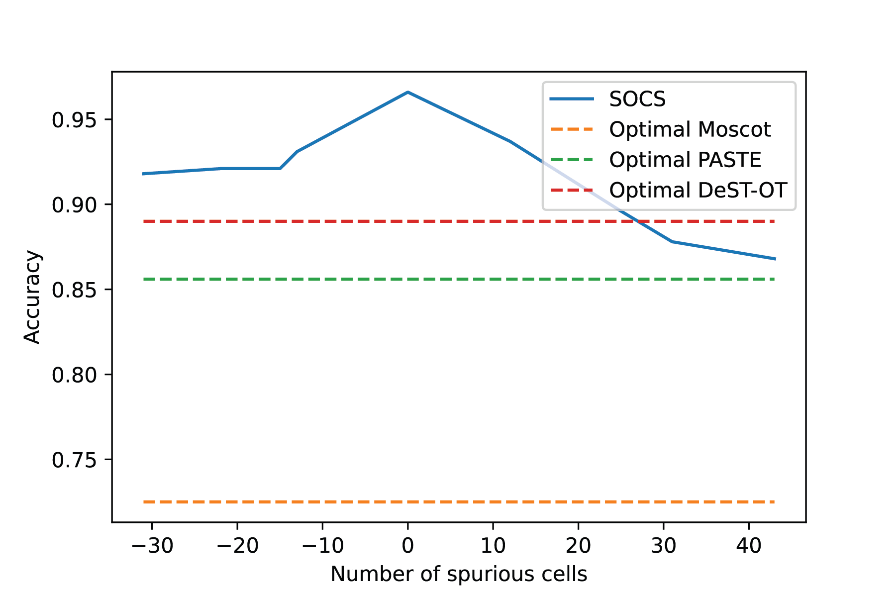

**Supplemental Figure 1 Dependence of mapping accuracy on segmentation:** Proportion of cells in the synthetic dataset mapping to the ground-truth descendant, varying based on the number of cells spuriously added (or removed from) the spatially coherent structure (with 282 cells in the ground-truth structure). Optimal accuracy of highly-performing alternative approaches for comparison.

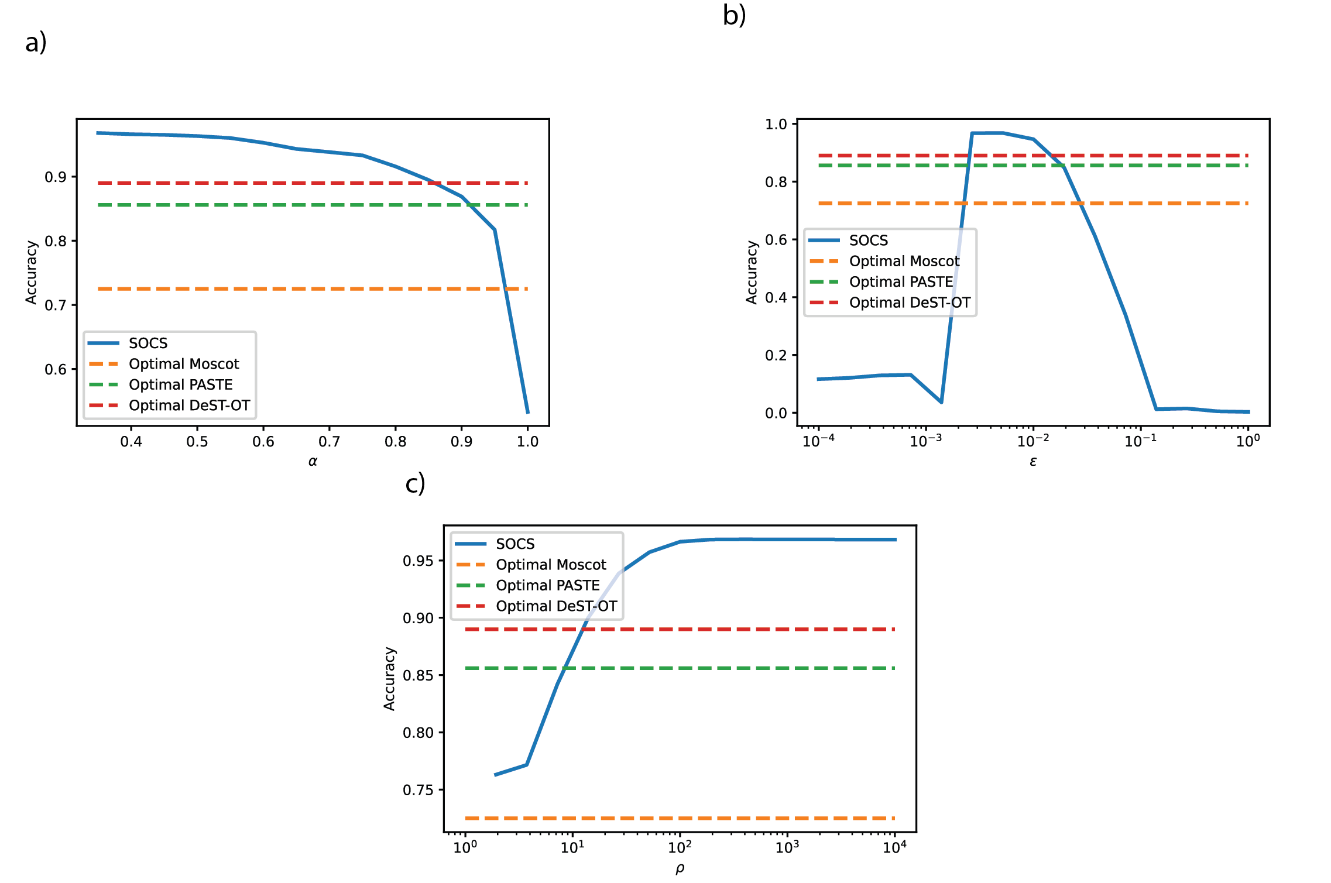

**Supplemental Figure 2 Dependence of mapping accuracy on parameter settings:** Proportion of cells in the synthetic dataset mapping to the ground-truth descendant, varying based on the tuning of individual parameters (with other parameters held constant at optimal values). a) varying $\alpha$, b) varying $\epsilon$, c) varying $\rho$ (we set $\rho_{1}=\rho_{2}=\rho$). Optimal accuracy of highly-performing alternative approaches for comparison.

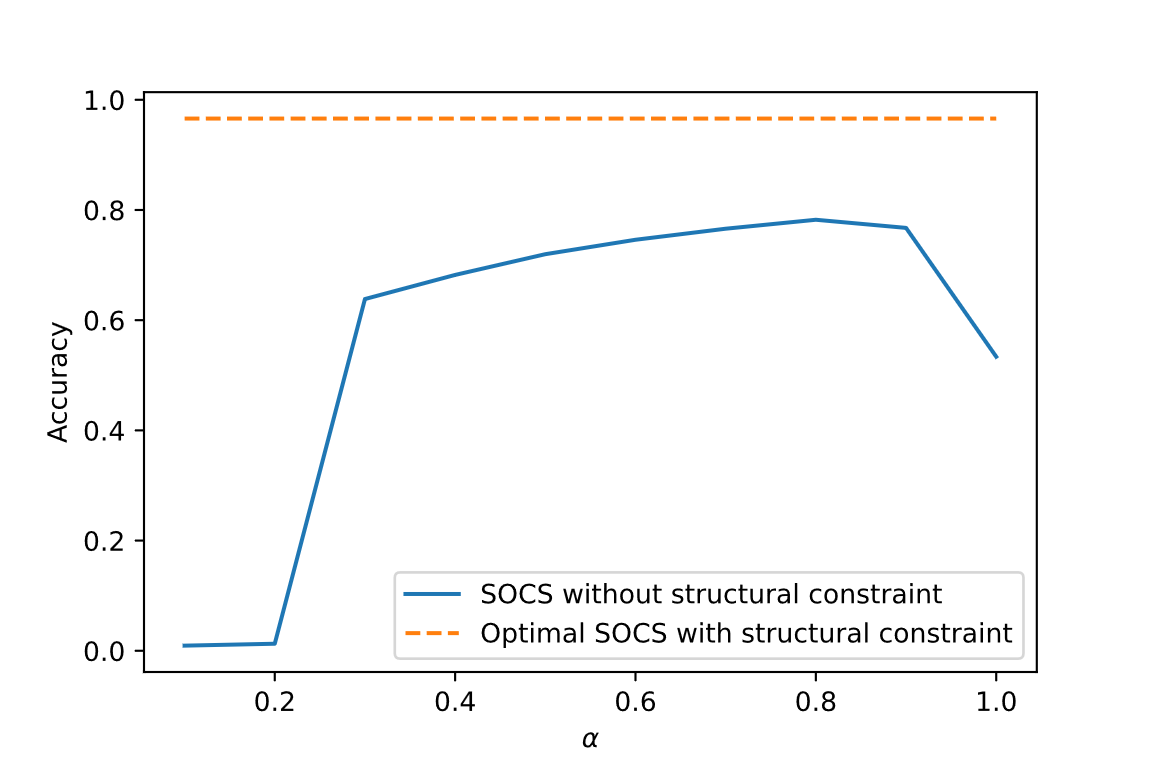

**Supplemental Figure 3 SOCS without structural constraint:** Proportion of cells in the synthetic dataset mapping to the ground-truth descendant in SOCS without the structural constraint, varying based on the weight assigned to spatial consistency. Optimal accuracy of SOCS with the structural constraint for comparison.

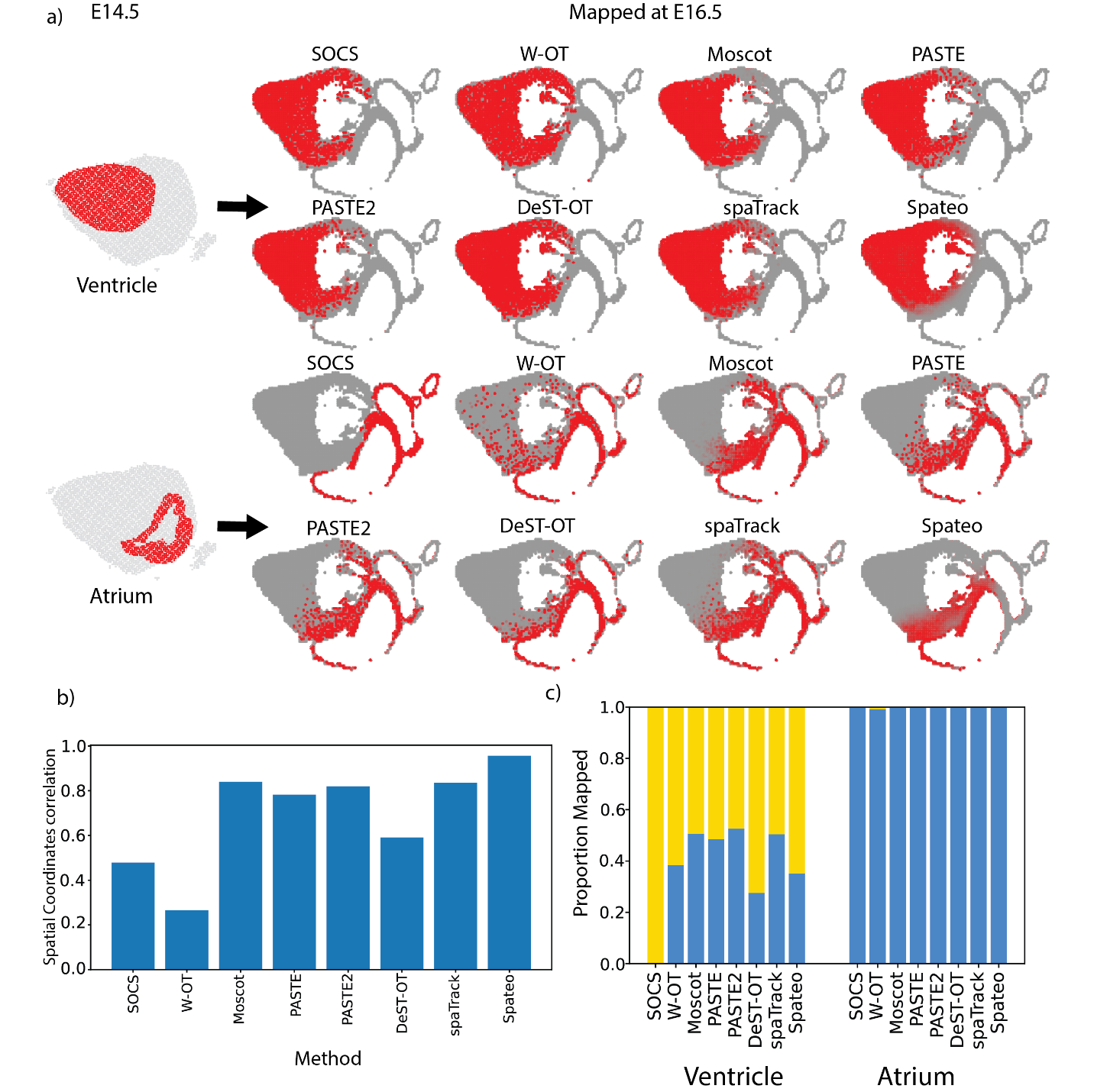

**Supplemental Figure 4 SOCS analysis of time-series Stereo-seq in developing mouse heart:** a) Cells from ventricle and atrium from mouse heart obtained at E14.5 mapped to E16.5 by SOCS and other methods. b) Scatter plots comparing pairwise spatial distances for each cell pair at E14.5 to their mapped pairwise distances at E16.5, by mapping method. c) Cell type mapping between mouse heart at E14.5 and E16.5, by mapping method.

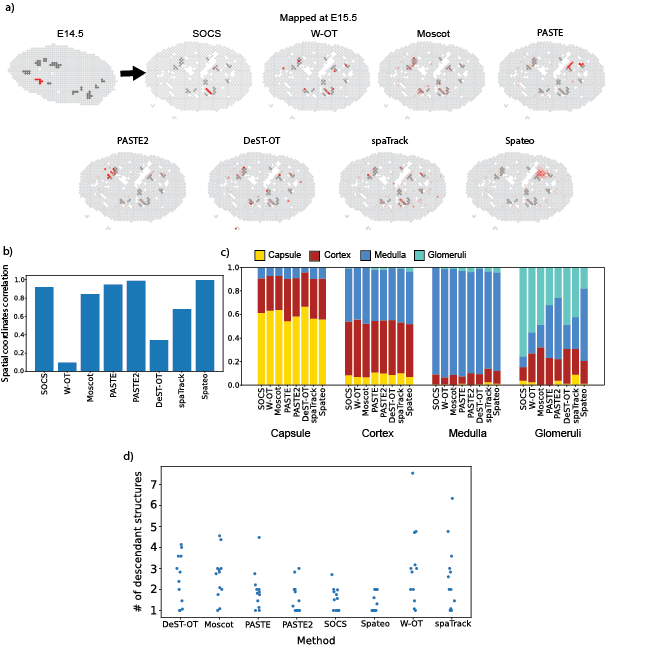

**Supplemental Figure 5 SOCS analysis of time-series Stereo-seq in developing mouse kidney:** a) Cells from a single glomerulus from mouse kidney obtained at E15.5 mapped to E16.5 by SOCS and other methods. b) Scatter plots comparing pairwise spatial distances for each cell pair at E15.5 to their mapped pairwise distances at E16.5, by mapping method. c) Cell type mapping between mouse kidney at E15.5 and E16.5, by mapping method. SOCS exhibits substantially stronger mapping from glomeruli to glomeruli than other methods. d) Strip plot showing effective number of descendant glomeruli at E16.5 for each glomerulus at E15.5 ($n=12)$, by mapping method.

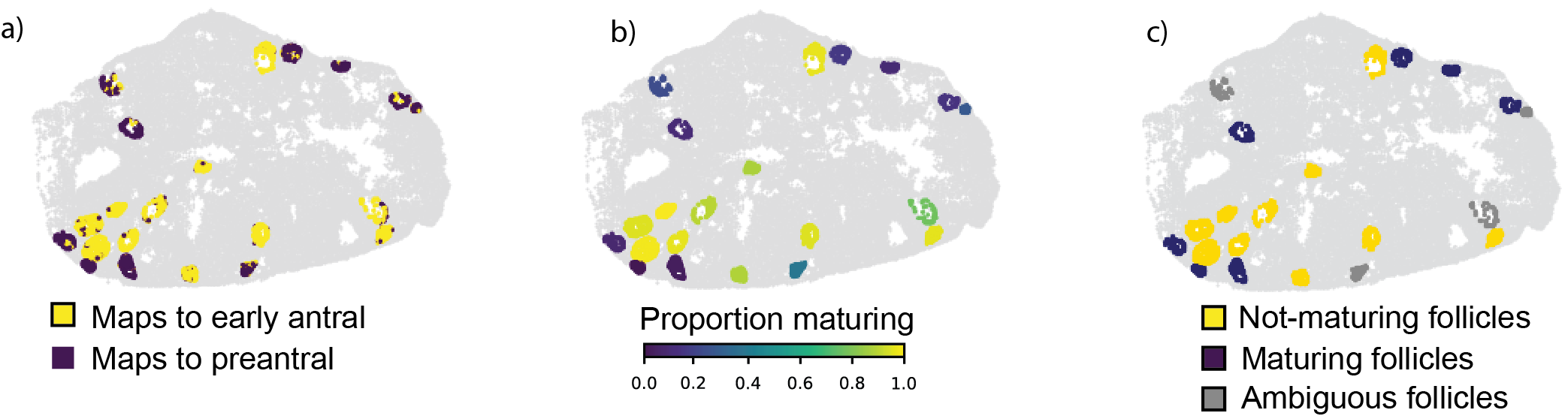

**Supplemental Figure 6 Categorizing maturing follicles:** a) Preantral follicle cells labeled by mapping to preantral or early antral follicle cells. b) Preantral follicles, colored by proportion of their cells mapping to early antral follicles. c) Follicles with 80% or more of cells mapping to early antral follicles labeled as “maturing,” follicles with 80% of more of cells mapping to preantral follicles labeled as “not maturing.”

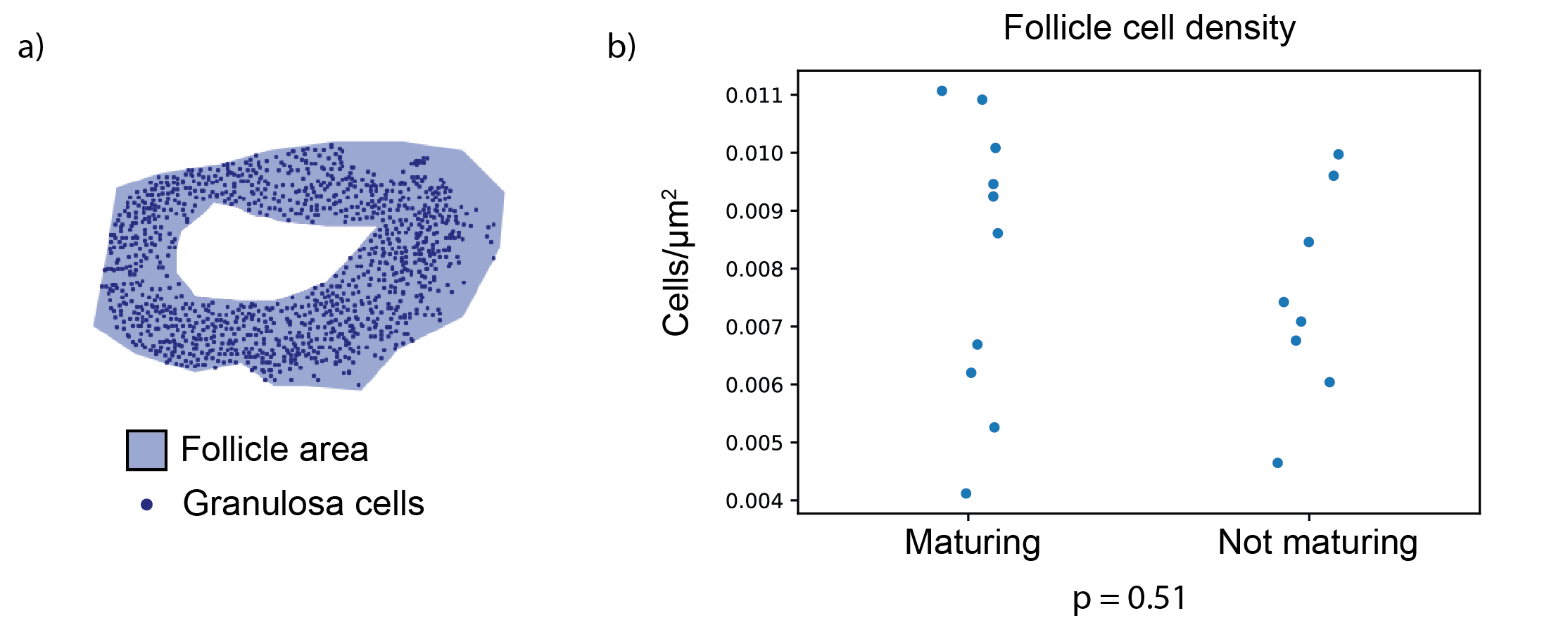

**Supplemental Figure 7 Follicle cell density comparison:** a) Computation of follicle density: the number of granulosa cells in the follicle is divided by the segmented follicle area. b) Comparison of cell density of maturing and not-maturing preantral follicles.

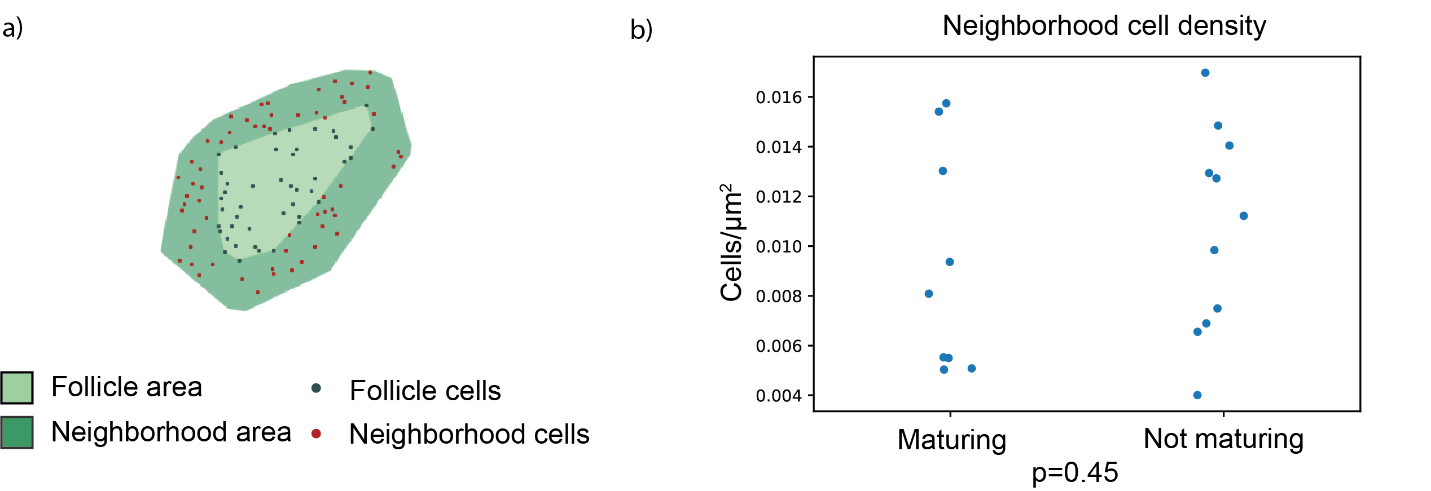

**Supplemental Figure 8 Follicle environmental density:** a) Computation of follicle neighborhood density: the number of cells in the follicle neighborhood is divided by the area of the neighborhood (obtained from the convex hull of the neighborhood cells). b) Comparison of neighborhood density of maturing and not-maturing preantral follicles

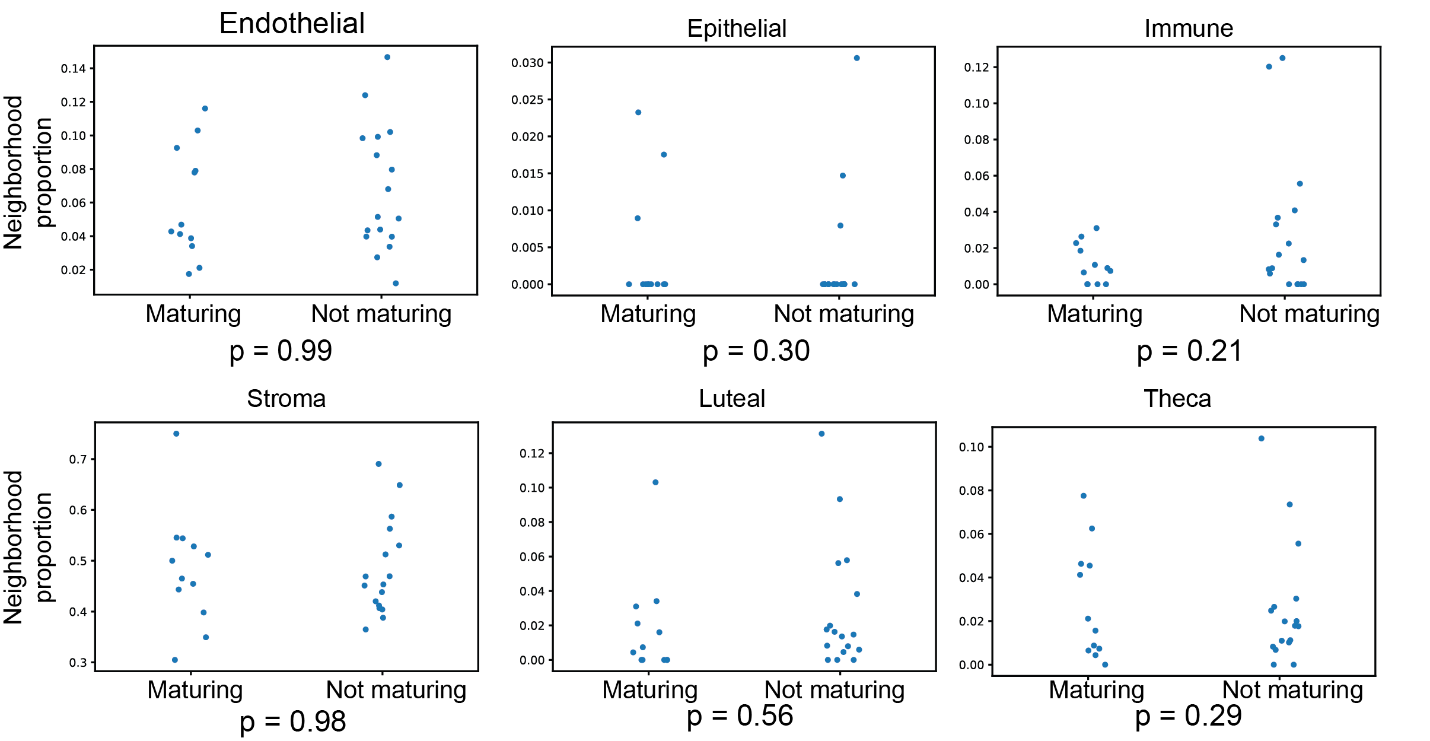

**Supplemental Figure 9:** For each major cell type, comparison of follicle neighborhood proportion in maturing and not-maturing preantral follicles.

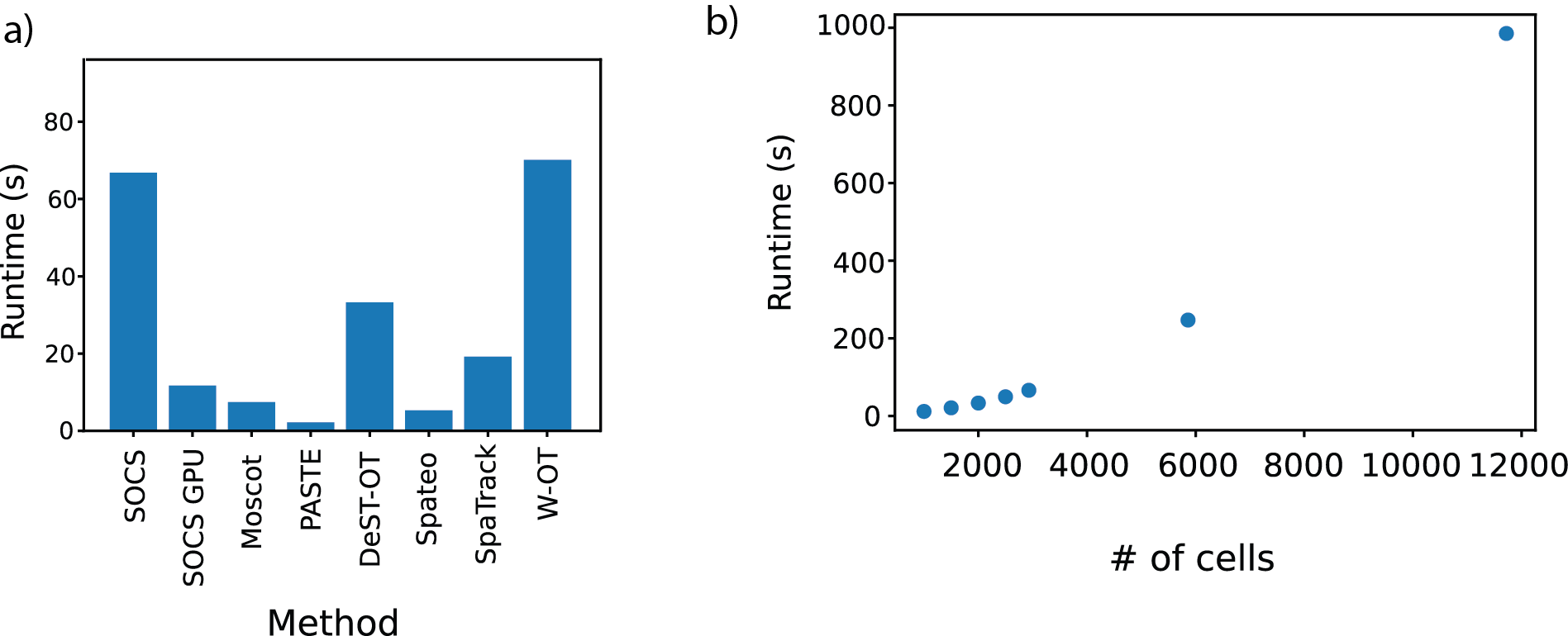

**Supplemental Figure 10 Runtime comparisons:** a) Time in seconds for each method to process two synthetic datasets composed of 1000 cells each. b) Time in seconds for SOCS GPU to process two synthetic datasets composed of varying numbers of cells.

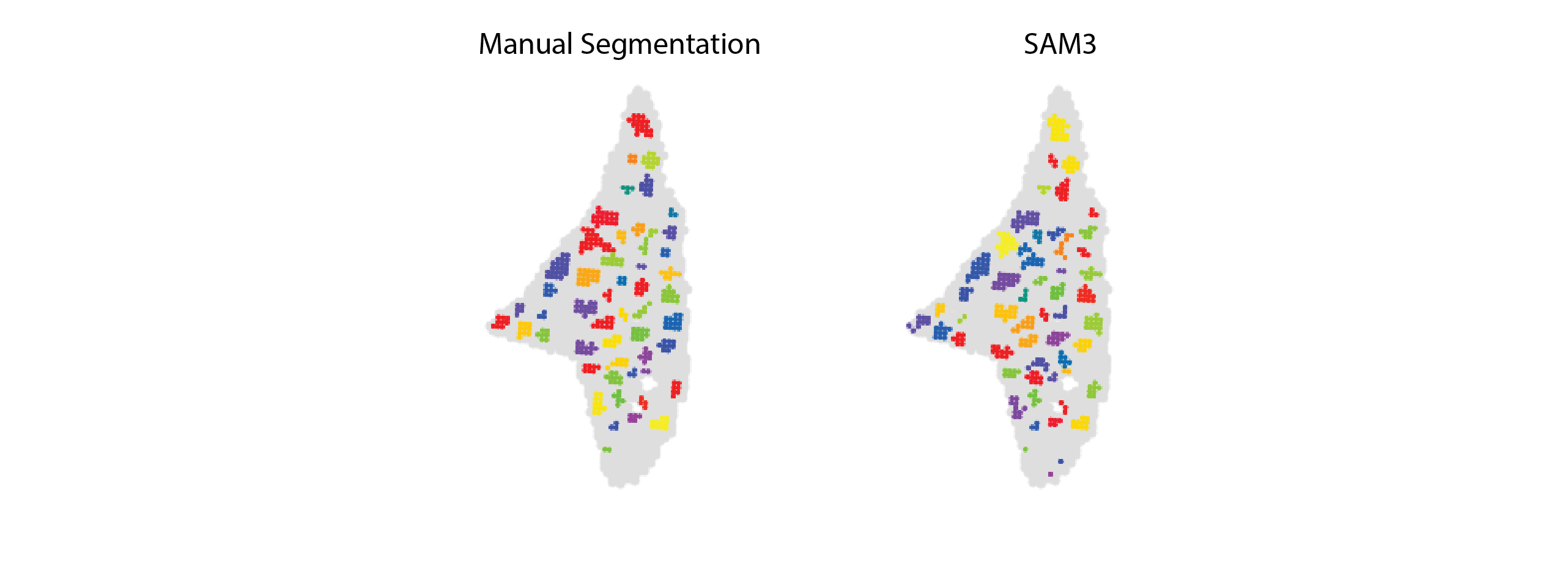

**Supplemental Figure 11 Automated segmentation with SAM3:** Comparison of structure segmentation of a slice of MOSTA lung organogenesis data, by manual segmentation (left) and SAM3 (right).

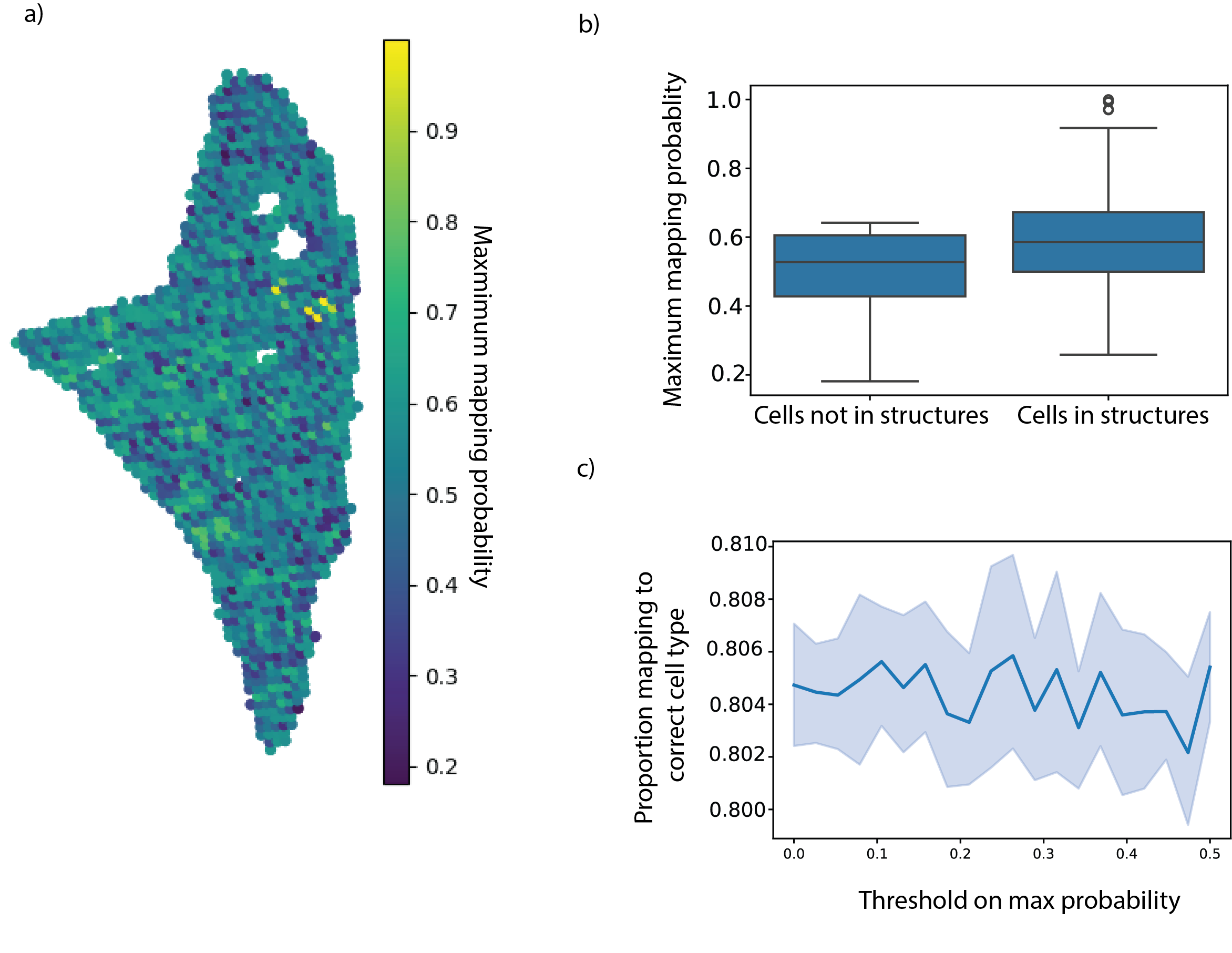

**Supplemental Figure 12 Confidence scoring based on mapping probability** a) spatial distribution of maximum mapping probability in the $t_{1}$ lung organogenesis data. b) Structure-associated cells tend to map with higher probability (center line is median, boxes represent interquartile range, whiskers extend to maximum and minimum elements excluding outliers). c) Filtering out cells which map with lower probability does not change the proportion of cells mapping to the correct cell type.
